# Isoflurane Anesthesia under Hyperoxic Conditions Shapes Cortical Hemodynamics and Photothrombotic Injury Severity in Mice: A Wide-Field Optical Imaging Study

**DOI:** 10.64898/2026.11.27.769604

**Authors:** N.V Lizunova, E.N. Kislukhina, V.I Dubas, V.I. Alekseev, A.M. Surin, Z.V. Bakaeva

## Abstract

**Background/Objectives:** Anesthesia can alter cortical hemodynamics and the severity of experimental brain injury. We investigated how isoflurane under hyperoxic conditions affects these.

**Methods:** Wide-field optical imaging (WFOI) in light-scattering mode (505 and 656 nm) through a chronic cranial window was used to assess hemoglobin dynamics in awake (n = 9) and isoflurane-anesthetized (n = 7) mice with intact cortices. Injury was assessed over 3 days after awake photothrombosis (Awake PT, n = 5), anesthetized photothrombosis (1.5% isoflurane/98.5% oxygen; IsO_2_ PT, n = 7), or laser irradiation without effective intravenous photosensitizer delivery (post hoc Laser group, n = 6). Assessments included FITC-dextran imaging of microcirculation and blood–brain barrier permeability, sensorimotor and biochemical tests, and 2,3,5-triphenyltetrazolium chloride (TTC) staining. Anesthesia induction was studied separately (n = 6).

**Results:** Anesthesia increased relative power at 0.05–0.10 Hz (p = 0.022) and primary motor cortex homotopic connectivity (p = 0.010), while reducing retrosplenial– sensorimotor correlations (p = 0.027). Anesthesia induction increased cortical blood oxygenation and total hemoglobin concentration. Mean lesion areas, early homotopic connectivity loss, day-1 forelimb asymmetry, and day-3 serum neuron-specific enolase increases followed a severity gradient (Laser < Awake PT < IsO_2_ PT). WFOI-derived lesion areas across the three groups at 10 min after irradiation were 2.03±0.99, 3.54±1.58, and 6.24±1.36 mm² (overall p < 0.001); day-3 TTC-negative areas were 1.60±0.73, 4.81±2.03, and 10.75±1.61 mm², respectively. IsO_2_ PT showed greater microvascular blockage, blood– brain barrier leakage, and approximately twofold longer hemodynamic responses to spreading depolarizations (p = 0.017) than Awake PT.

**Conclusions:** Isoflurane anesthesia under hyperoxic conditions affects baseline cortical activity and exacerbates photothrombotic injury, potentially through enhanced oxygen-dependent photochemistry, emphasizing the importance of anesthetic regimens in experimental models.

## Introduction

Acquired brain injuries, including ischemic stroke and traumatic brain injury, represent a major medical and socioeconomic burden (Feigin et al., 2024; Feigin et al., 2025). Experimental models that reproduce key pathological processes and allow their consequences to be assessed are essential for investigating the mechanisms of brain injury. Many studies in laboratory animals, including those involving the induction of ischemic injury, are performed under anesthesia. However, anesthesia is not a physiologically neutral condition and may affect both the recorded brain activity and the development of injury itself.

Anesthesia has been shown to alter the spatiotemporal organization of spontaneous cortical activity and functional interactions between brain regions (Mohajerani et al., 2010; Wright et al., 2017; West et al., 2022). Anesthetics may also influence the severity of experimental ischemic injury. Neuroprotective effects of isoflurane have been reported in several models (Archer et al., 2017), including photothrombosis in mice (Seto et al., 2014). Nevertheless, the magnitude and direction of these effects depend on the experimental conditions; for example, after transient middle cerebral artery occlusion, injury outcomes were less severe under propofol anesthesia than under isoflurane anesthesia (Franx et al., 2024).

The composition of the inhaled gas mixture and ventilation parameters are also important (Liu et al., 2009). In particular, the use of oxygen as the carrier gas for isoflurane, often supplied as 100% O₂, remains a common practice in experimental photothrombosis protocols (Zheng et al., 2010; Lechtenberg et al., 2019; Collyer et al., 2023; Boyle et al., 2026). On the one hand, hyperoxia may improve penumbral tissue oxygenation during focal cerebral ischemia, although its effects depend on the composition of the administered gas mixture (Liu et al., 2006). On the other hand, elevated inspired oxygen concentrations may cause cerebral vasoconstriction (Floyd et al., 2003), accelerate thrombus formation during Rose Bengal-mediated photochemical vascular injury in retinal vessels (Wilson and Hatchell, 1991), and alter gas exchange, including by promoting atelectasis (Wilding et al., 2017). During spontaneous breathing under anesthesia, reduced ventilation may lead to CO₂ accumulation and cerebral vasodilation (Sicard and Duong, 2005). Other components of inhaled gas mixtures, such as nitrous oxide, are not pharmacologically inert either (Baughman et al., 1989; Yokoo et al., 2004; Haelewyn et al., 2008). Thus, the effects of an anesthetic on injury development must be considered in the context of the inhaled gas composition and the animal’s physiological state. Although isoflurane has been shown to influence early hemodynamic changes and lesion development during photothrombotic stroke induction (Lu et al., 2017), the combined effects of isoflurane anesthesia and elevated inspired oxygen concentrations under these conditions remain insufficiently characterized.

Among the acute consequences of ischemic injury, blood–brain barrier (BBB) disruption and spreading depolarizations (SDs) are of particular importance. Increased BBB permeability contributes to neuroinflammation and vasogenic edema and is associated with a risk of hemorrhagic transformation (Hoffmann et al., 2012; Blase et al., 2025). Spreading depolarizations can exacerbate the mismatch between metabolic supply and demand in ischemic tissue and may promote lesion expansion, including through spreading ischemia (Shin et al., 2006; Nakamura et al., 2010; von Bornstädt et al., 2015; Ayata and Lauritzen, 2015). The characteristics of SDs also depend on anesthetic conditions (Kudo et al., 2008; Klass et al., 2018). Assessment of early vascular and functional disturbances is therefore important for understanding how injury-induction conditions influence injury severity.

Wide-field optical imaging (WFOI) can be used to investigate these processes, allowing changes over a substantial portion of the cortical surface to be recorded repeatedly in the same animal, including during wakefulness (Vanni and Murphy, 2014; Cardin et al., 2020; Ma et al., 2016a; Kislukhina et al., 2026). In optical intrinsic signal imaging (OISI) mode, the technique enables estimation of changes in the concentrations of oxygenated (HbO), deoxygenated (HHb), and total hemoglobin (HbT) without genetically encoded indicators (White et al., 2011; Kura et al., 2018). Like functional magnetic resonance imaging (fMRI) and functional near-infrared spectroscopy (fNIRS), WFOI measures hemodynamic changes coupled to neuronal activity through neurovascular coupling (Ferrari and Quaresima, 2012; Hillman, 2014; Yatsyk et al., 2023). Repeated measurements enable comparison of cortical status before and after an intervention and assessment of functional connectivity, lesion dynamics, and early hemodynamic disturbances.

The present study had two primary objectives: first, to investigate the effects of isoflurane anesthesia with oxygen as the carrier gas on spontaneous hemodynamic activity and functional connectivity in the intact mouse cortex, as measured by WFOI. And second, to determine how these experimental conditions, applied during photothrombosis induction, influence the initial extent, subsequent development, and functional consequences of cortical injury. To address the latter objective, we compared early and delayed structural, vascular, and functional changes, including lesion-area dynamics, impaired microcirculation and BBB permeability, spreading depolarizations, and motor and biochemical outcomes.

## Materials and Methods

### Animals, Ethics, and Study Design

The main experiment initially included 20 male C57BL/6 mice, and an additional 6 mice were used in a separate experiment. Animals were approximately 2 months old and weighed 25–30 g. Before the experimental procedures, the mice were acclimatized for 14 days and housed at 22 °C under a 12-h light/dark cycle with ad libitum access to food and water. Animals were group-housed (5–6 per cage) before surgery and individually housed after preparation of the chronic cranial window. All procedures complied with Directive 2010/63/EU and were approved by the local Ethics Committee of the National Medical Research Center for Children’s Health (Protocol No. 1, 30 January 2018).

The study used a longitudinal design, with the individual animal serving as both the experimental and statistical unit. No a priori sample-size calculation was performed. Before the main longitudinal experiment, animals were randomly allocated to photothrombosis induction under awake conditions or isoflurane anesthesia. Before Rose Bengal administration, baseline resting-state WFOI recordings were obtained for 6 min in awake or isoflurane-anesthetized mice, according to their assigned experimental conditions. Following recording-quality assessment, independent groups of Awake (n = 9) and Isoflurane (n = 7) animals were included in the analysis of spontaneous hemodynamic activity in the intact cortex. Two mice were subsequently excluded from injury-outcome analyses because of deviations from the injury-induction protocol. In five mice, the full intended dose of Rose Bengal (10 mg/kg) was administered through an incorrectly positioned or nonpatent catheter, resulting in presumed subcutaneous rather than intravenous delivery. One additional mouse (m18) received an equivalent volume of saline instead of Rose Bengal through a patent catheter. These six animals constituted a post hoc exploratory comparator group (Laser). The final injury-outcome groups were Awake PT (n = 5), IsO_2_ PT (n = 7), and Laser (n = 6). The Laser group included animals exposed to laser irradiation either while awake (n = 4) or under isoflurane anesthesia (n = 2).

Two mice in the IsO_2_ PT group died within the first 24 h after injury; their data were used only for available acute outcomes and were excluded from subsequent longitudinal analyses. Individual measurements were excluded in cases of technical failure, insufficient recording quality, or unusable biological samples, without excluding the remaining data from the same animal. The actual number of animals is reported for each analysis. Formal blinding was not applied; however, group assignments were not specifically used during WFOI data processing.

The experimental design and sequence of procedures are shown in Figure 1.

**Figure 1.**
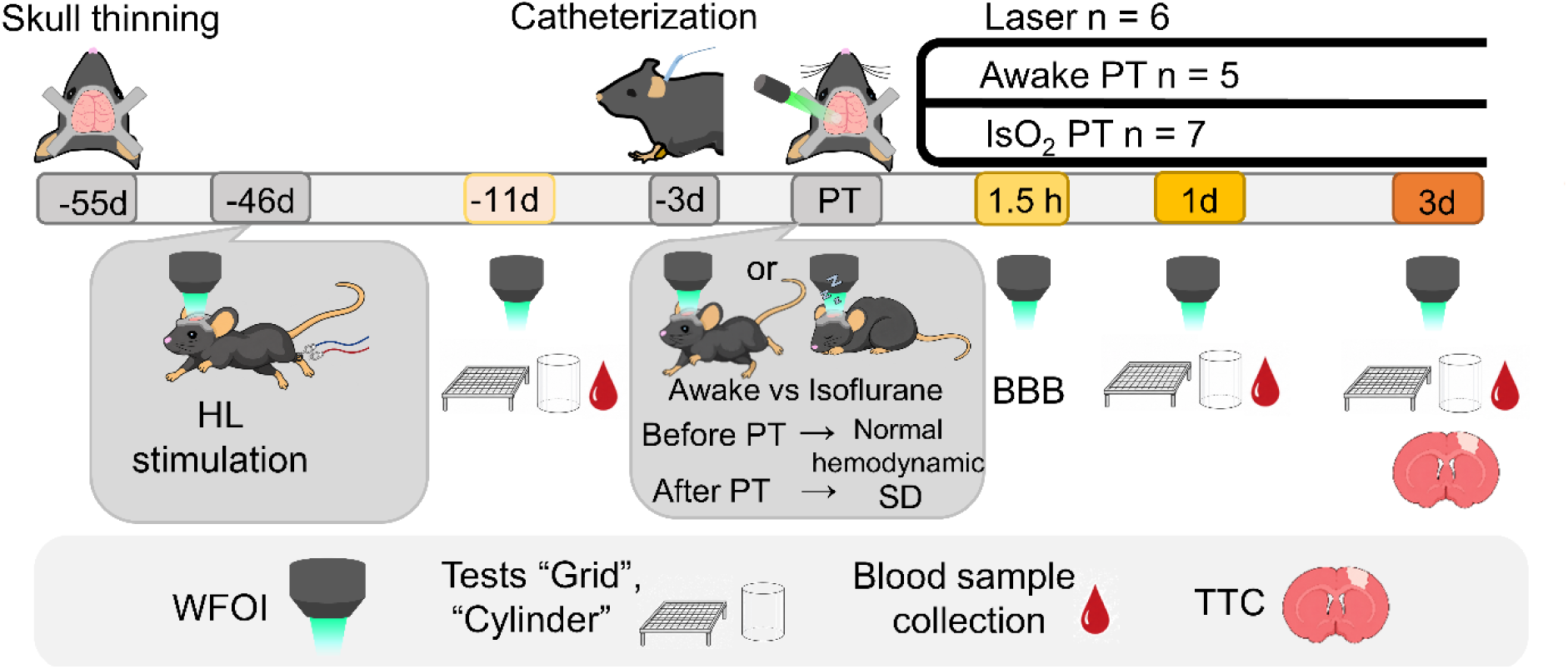
Experimental design. The main experimental procedures and time points for assessments before and after induction of focal cortical injury are shown. WFOI, wide-field optical imaging; PT, photothrombosis; SD, spreading depolarization; BBB, blood–brain barrier; TTC, 2,3,5-triphenyltetrazolium chloride; HL, hindlimb.

### Chronic Cranial Window Surgery

At 55 (41–62) days before injury induction (here and throughout this section, median (range)), animals underwent chronic cranial window surgery according to a previously described protocol (Kislukhina et al., 2026), with modifications. After the soft tissues over the dorsal skull were removed, the skull bone was thinned, covered with transparent polymer material, and fitted with a titanium head holder. During surgery, ring electrodes made of 0.5-mm-diameter steel wire were attached around the Achilles tendon for subsequent somatosensory stimulation. Unlike the previously published protocol, the procedure was performed under injectable anesthesia with tiletamine/zolazepam (Zoletil 100, Virbac, France) and xylazine (Xyla, Interchemie werken “De Adelaar” B.V., the Netherlands), at doses of 100 and 8 mg/kg, respectively, administered intraperitoneally. In addition, the titanium holder had a narrower, elongated shape along the rostrocaudal axis. The holder was secured with dental acrylic without a plastic adapter. Body temperature was maintained at 37 °C, and anesthetic depth was monitored by the absence of the pedal withdrawal reflex. After surgery, each mouse was placed in an individual cage and monitored until full recovery from anesthesia. For 3 days after surgery, animals received ketoprofen (Sandoz, Slovenia; 8.5 mg/kg, injection volume 5 µL/g body weight) once daily by subcutaneous injection, and the electrode implantation site was treated with 10% povidone–iodine (Betadine, EGIS Pharmaceuticals, Hungary).

### Wide-Field Optical Imaging

Cortical hemodynamic activity was recorded by WFOI in light-scattering mode. The optical system comprised LED light sources at 505 nm (BLS-LCS-0505-12-22; Mightex, USA) and 656 nm (BLS-LCS-0656-12-22; Mightex, USA), a 4× epi-objective (P/N OBJ-MAO-04; Mightex, USA), and a high-sensitivity camera (CXE-B013-U; Mightex, USA). Light from the LEDs was directed to the brain surface through ET505/20x (Chroma Technology Corp., VT, USA) and AT656/30m (Chroma Technology Corp., VT, USA) optical filters, a combined beam splitter (LCS-BC25-0515; Mightex, CA, USA), and a custom-built side-illumination system. Diffusely scattered light was collected in an upright microscope configuration through a 59009bs beam splitter and a 59026m emission filter (Chroma Technology Corp., VT, USA).

During the main hemodynamic recordings, illumination at 505 and 656 nm was alternated sequentially. The total acquisition rate was 33.3 frames/s, corresponding to an effective rate of 16.67 frames/s at each wavelength. Raw data were stored as 8-bit BMP images (464 × 344 pixels).

During awake recordings, the head was fixed relative to the optical system while the animal could move freely on an air-lifted platform (Neurotar, Finland). Gradual habituation to WFOI conditions followed a previously published protocol (Kislukhina et al., 2026), except that habituation began 7 days after surgery.

### Somatosensory Stimulation

At 8 (7–10) days after surgery, a 12-min recording with electrical hindlimb stimulation was acquired in awake animals. Stimulation was delivered via implanted electrodes using a TTL-controlled JDS6600-60M stimulator (Juntek, China) with adjustable voltage. Each train comprised four pulses delivered at 1 Hz, with a duration of 400 ms per pulse; trains were presented at 360, 450, 540, and 630 s after recording onset. The actual current was 150–350 µA and was monitored with a Hantek365A digital ammeter. Stimulation was used to functionally localize the cortical hindlimb representation and subsequently register the anatomical atlas for each animal.

### Jugular Vein Catheterization

At 3 (2–4) days before injury induction, a chronic polyethylene catheter (TPM-E-0.3 × 0.15; Medsila JSC, Russia) was implanted into the internal jugular vein for administration of Rose Bengal and fluorescein isothiocyanate (FITC)-labeled dextran. Surgery was performed under tiletamine/zolazepam and xylazine anesthesia (100 and 8 mg/kg, respectively). The catheter was filled with heparin (50 IU/mL; 5 µL/mouse; Belmedpreparaty, Belarus). Patency was assessed by resistance during injection and subsequently confirmed by visualizing the entry of fluorescent dextran into the bloodstream.

### Induction of Focal Cortical Injury by Photothrombosis

Focal cortical injury was induced by photothrombosis based on a previously described protocol (Galkov et al., 2020). Rose Bengal (198250-5G; Sigma-Aldrich) was administered through the jugular catheter at 10 mg/kg (200 µL per 30 g body weight). Five minutes later, the right somatosensory cortex was irradiated for 5 min with a 532-nm laser (KSK, China); nominal power was approximately 1000 mW, and the beam diameter was 1.5 mm. The center of irradiation was located at AP = −1 mm and ML = −2 mm relative to bregma (as defined in the original protocol, the negative directions indicated caudal and rightward displacement, respectively).

Animals in the Awake PT group were head-fixed in the WFOI setup without general anesthesia. For the IsO_2_ PT group, anesthesia was induced in an anesthesia chamber (4% isoflurane, 96% O₂, flow rate 1 L/min); animals were then transferred to the imaging setup, where anesthesia was maintained through a face mask (1.5% isoflurane, 98.5% O₂, flow rate 1 L/min). These concentrations represent the nominal vaporizer and flowmeter settings; the actual inspired gas composition was not independently measured.

Following a 6-min baseline resting-state WFOI recording in awake or isoflurane-anesthetized mice, Rose Bengal was administered, and the catheter was flushed with heparin (10 IU/mL; 10 µL/mouse; Belmedpreparaty, Belarus). 5 min were allowed for dye distribution, followed by 5 min of laser irradiation. A 12-min recording to assess acute hemodynamic and vascular changes was initiated approximately 3 min after irradiation ended, under the same anesthetic or awake conditions.

In five animals subsequently included in the Laser group, intravenous Rose Bengal delivery was unsuccessful, as indicated by the absence of FITC-dextran entry into the bloodstream during the subsequent vascular assessment. In the remaining Laser animal (m18), saline was administered instead of Rose Bengal. Laser irradiation was performed according to the same protocol in all six animals.

### Longitudinal Follow-Up

At 11.0 (9.0–12.0) days before injury, baseline motor testing, blood sampling, and a 6-min WFOI recording of spontaneous cortical hemodynamic activity were performed in awake animals. After injury, 6-min recordings of spontaneous cortical hemodynamics were obtained at 1.5 h, 24 h, and 72 h. Motor testing and blood sampling were repeated at 24 and 72 h after injury.

### Assessment of Blood–Brain Barrier Permeability and Capillary Perfusion

To assess vascular perfusion and BBB permeability, fluorescently labeled 10-kDa FITC-dextran (FD10S-100MG, Sigma-Aldrich, USA) was administered intravenously through the jugular catheter 1.5 h after photothrombosis, at a dose of 30 µg/g body weight and an injection volume of 3 µL/g. Administration was recorded by WFOI in fluorescence mode using excitation at approximately 470 nm and emission detection at approximately 525 nm. Chroma filter set #49002 was used, including ET470/40x and ET525/50m excitation and emission filters, respectively, and a T495lpxr dichroic mirror. First-pass dextran dynamics were recorded for 1 min at 33.3 frames/s.

The area lacking capillary-bed filling was determined from the dynamic first-pass dextran recordings. The area of dextran extravasation into tissue was assessed from images obtained 1 min after administration. To evaluate extravasation, several spatially offset images were registered and combined into a maximum-intensity projection. Masks delineating the regions of capillary perfusion blockage and extravasation were created manually in ImageJ-Fiji (v1.54f). Before delineation, mean image intensity was normalized across animals using three regions of interest (ROIs) in the intact hemisphere.

In five animals in the Laser group, intravenous FITC-dextran administration was unsuccessful because of defective catheters. In the remaining Laser animal (m18), which had received saline instead of Rose Bengal, FITC-dextran imaging was used only as an illustrative reference. The Laser group was not included in statistical comparisons of capillary perfusion impairment or dextran extravasation.

### Blood Sampling and Enzyme-Linked Immunosorbent Assay

Blood was collected by puncture of the submandibular vein before injury and at 24 and 72 h afterward. After clot formation for 2 h at room temperature, samples were centrifuged at 1000 × g and 4–8 °C for 20 min; serum was stored at −70 °C.

Serum neuron-specific enolase (NSE) concentration was measured by enzyme-linked immunosorbent assay (ELISA) using the ELISA Kit for Enolase, Neuron Specific (Cloud-Clone Corp., Wuhan, China; SEA537Mu), following the manufacturer’s instructions. The detection range was 78– 5000 pg/mL, and sensitivity was <32 pg/mL. A four-parameter logistic calibration model (4PL; R² = 0.999) was used. S100β was measured by ELISA using the ELISA Kit for S100 Calcium Binding Protein B (Cloud-Clone Corp., Wuhan, China; SEA567Mu), following the manufacturer’s instructions. The detection range was 31.2–2000 pg/mL, and sensitivity was <12.5 pg/mL. A four-parameter logistic calibration model (4PL; R² = 1) was used. Serum samples were analyzed at dilutions ranging from 1:4 to 1:2. Measurements were performed in duplicate for day 3 samples when the available sample volume was sufficient.

### Behavioral Tests

Sensorimotor deficits were assessed using the Grid and Cylinder tests before catheterization and at 24 and 72 h after injury (Baskin et al., 2003; Brooks and Dunnett, 2009). Before testing, mice were acclimated to a darkened room for 30 min.

In the Grid test, the animal moved freely across a horizontal grid with 1-cm openings until 100 steps had been completed. A limb entering a grid opening was scored as an error; the number of errors made by the left limbs per 100 steps was analyzed.

In the Cylinder test, video recordings of 15 rearings were used to determine the duration of contact of the left and right forelimbs with the cylinder wall. The asymmetry index was calculated as the ratio of total left-forelimb support time to total support time involving both forelimbs. Results were expressed as a percentage.

### Experiment Termination and TTC Staining

At 72 h, animals were euthanized by an overdose of tiletamine/zolazepam and xylazine followed by decapitation. Brains were sectioned into 2-mm-thick coronal slices, which were incubated in 0.5% 2,3,5-triphenyltetrazolium chloride (TTC) at 37 °C for 15 min and turned over after 7.5 min. After staining, the slices were fixed in 10% formalin, photographed, and analyzed using ImageJ-Fiji (v1.54f). Images were first scale-calibrated; the lesion and total hemispheric area were then delineated using the Freehand Selections tool, and areas were measured using the Measure command. For quantitative analysis, the coronal section showing the largest TTC-negative lesion area was selected for each animal.

### Hemodynamic Changes during Isoflurane Anesthesia Induction

The additional experiment was conducted in six male C57BL/6 mice. Initially, the animals were awake in the WFOI setup. A face mask delivering 100% oxygen was placed near the animal’s snout, and continuous recording was initiated. Seven seconds after recording onset, delivery of 4% isoflurane was started; after a further 45 s, the concentration was reduced to 1.5%. Total recording duration was 12 min. Imaging was performed at 505 and 656 nm with an effective acquisition rate of 16.67 Hz. Changes in HbO, HHb, and HbT concentrations were quantified as the difference between the mean values during the final 60 s and the first 7 s of the recording.

### Wide-Field Optical Imaging Data Processing

Image processing and quantitative analysis were performed using custom code developed for this project in Python 3.11.15 with NumPy 2.4.6, SciPy 1.17.1, pandas 3.0.3, scikit-image 0.26.0, xarray 2026.4.0, and OpenCV 5.0.0. Pixel classification for automated mask generation was performed in Ilastik (v 1.4.1). Visual inspection, manual mask correction, registration quality control, and anatomical annotation were performed using interactive interfaces based on napari (v 0.7.0).

### Geometric Preprocessing and Mask Generation

Raw images were rotated by −90°. For hemodynamic-signal calculation, the linear image dimensions were reduced to 30% of the originals.

Brain and technical-artifact masks were derived from Ilastik pixel-classification probability maps. The brain mask was defined on 656-nm images using a probability threshold of 0.60. Regions containing bregma, glare, contamination, and vascular structures were masked separately in the 505-and 656-nm images. Automatically generated brain masks and bregma locations were inspected visually and corrected manually when necessary.

### Intersession Registration

For each mouse, images acquired on different experimental days were registered to an individual reference using the 505-nm channel. Intensity was normalized to the 1st and 99th percentiles; low-frequency spatial background was removed using a Gaussian filter (σ = 12 pixels), and vascular patterns were enhanced using a Sobel operator. Initial alignment was based on bregma, followed by optimization of rigid rotation about bregma over ±15° in 0.1° increments to maximize the overlap between Sobel maps. All registrations were visually inspected, with translation and rotation adjusted manually if needed.

### Calculation of Hemodynamic Signals

Wide-field light-scattering images acquired at 505 and 656 nm were converted into changes in oxyhemoglobin (ΔHbO), deoxyhemoglobin (ΔHHb), and total hemoglobin (ΔHbT) concentrations using the modified Beer–Lambert law and two-wavelength spectroscopic decomposition, as described by Ma et al. (2016a). At each wavelength, the intensity was corrected for slow illumination drift by dividing by the mean signal from a background region outside the brain; each pixel was then normalized to its mean intensity over the entire recording:

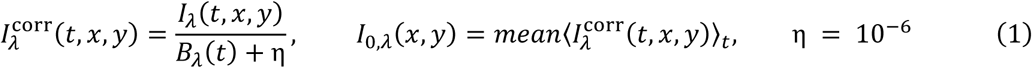

The change in optical density and the corresponding change in the absorption coefficient were calculated as follows:

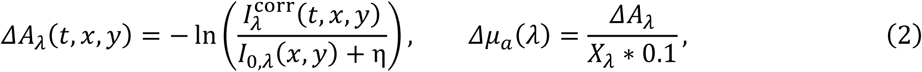

where *B_λ_*(*t*) is the mean background signal, *I*_0,_*_λ_* is the mean pixel intensity over the recording, *ΔA_λ_* is the change in optical density, *X_λ_* is the effective optical path length (mm), and *Δμ_a_*(*λ*) is the change in the absorption coefficient. Effective optical path lengths were taken from Ma et al. (2016a).

For *λ*_1_ = 505 nm and *λ*_2_ = 656 nm, concentration changes were reconstructed using the extinction coefficients *ξ*_HbO_(*λ*) and *ξ*_HHb_(*λ*) as follows:

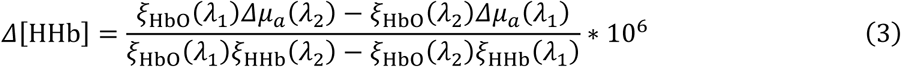

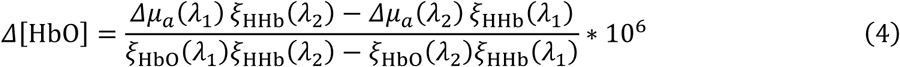

Total hemoglobin was calculated as follows:

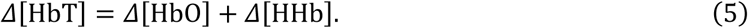

Tabulated base-10 molar extinction coefficients (ε) were obtained from the Oregon Medical Laser Center (Prahl, n.d.) and converted to natural-logarithm-based coefficients by multiplication by ln(10) (ξ).

Calculated concentrations were converted to µM. After spectroscopic decomposition, the global hemodynamic component was removed from recordings with sensory stimulation by subtracting the cortex-wide mean signal; no global signal regression was performed for the spectral or functional-connectivity analyses. Before final analysis, signals were additionally smoothed with a Gaussian filter (*σ_xy_* = 1 pixel, *σ_t_* = 1 frame).

### Behavioral Annotation and REST-Segment Selection

Periods of locomotion and quiet wakefulness were annotated manually from synchronized behavioral videos while simultaneously inspecting the hemodynamic signal. If video was unavailable, WFOI signal features established from recordings with video were used: the absence of prominent abrupt amplitude changes and a signal fluctuation range characteristic of periods without active locomotion. Continuous 60-s epochs of quiet wakefulness (REST) were used for the main analysis of spontaneous activity. Candidate epochs underwent additional visual quality control, and segments with prominent movement or technical artifacts were excluded. Under isoflurane anesthesia, the entire recording was divided into consecutive 60-s segments, all of which were included in the analysis. Multiple REST segments from the same recording were treated as repeated measurements of a single biological unit rather than independent observations.

### Registration to the Allen Mouse Brain Atlas

Individual images were registered to the Allen Mouse Brain Common Coordinate Framework CCFv3 (allen_mouse_25um, BrainGlobe) based on bregma position, the rostrocaudal axis, and localization of the hemodynamic response to hindlimb stimulation. The maximum-response region was identified manually in napari from data averaged across four stimulus presentations, with particular emphasis on the early, spatially localized component of the response. The optimization allowed the tilt of the dorsal atlas surface to vary from −10° to 20° and the scale factor from 0.80 to 1.40.

Regional analyses included MOp (primary motor cortex), MOs (secondary motor cortex), SS (somatosensory cortex), and RSP (retrosplenial cortex). To reduce uncertainty at region boundaries, a 2-pixel boundary strip in canonical atlas space, a ±12-pixel region along the midline, and a 20-pixel peripheral strip of the common technically accessible mask were excluded. Only ROIs with sufficient reproducible spatial coverage were included.

### Functional Connectivity Analysis

Functional connectivity (FC) was analyzed at frequencies below 0.40 Hz. This range was selected based on studies showing substantial correspondence between spontaneous hemodynamic and neuronal signals within this frequency range (Ma et al., 2016b; Wright et al., 2017). Connectivity was calculated separately for each accepted 60-s REST segment of the HbT signal. A linear trend was removed from each valid time series before applying a fourth-order Butterworth low-pass filter with a cutoff frequency of 0.40 Hz. Zero-phase filtering was implemented in the forward and reverse directions. To reduce edge effects, the first and last 2 s of each 60-s segment were removed after filtering. For pixelwise analysis, linear spatial resolution was further reduced by a factor of two while accounting for the valid-pixel mask.

Homotopic connectivity (HC) was defined as the Pearson correlation coefficient between the time series of spatially corresponding mirror-image pixels in the two hemispheres. The resulting correlation coefficients were transformed using Fisher’s z transformation:

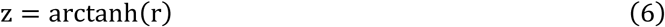

The z maps were calculated independently for each REST segment and subsequently averaged within a recording. The regional value was determined as the mean across valid homotopic pixel pairs within the corresponding ROI.

Global brain connectivity (GBC) for each pixel was defined as the mean Fisher-z-transformed correlation between its time series and those of all other valid pixels in the analyzed cortex. In the main longitudinal analysis after injury, GBC was assessed in the contralateral, morphologically intact hemisphere.

Interregional functional connectivity was assessed using Pearson correlations between regionally averaged HbT time series, obtained by averaging the signals from the left and right hemispheres. Correlation coefficients were Fisher-z-transformed and averaged at the animal level. Mean RSP connectivity was calculated for each animal by averaging Fisher z values for three region pairs (RSP–MOp, RSP–MOs, and RSP–SS). Only animals with valid data for all three pairs were included in this analysis.

### Spectral Analysis of Spontaneous Hemodynamic Activity

Spectral analysis was performed on HbT signals from accepted REST segments. For each ROI and hemisphere, a spatially averaged time series was derived from valid pixels. The linear trend was removed before spectral analysis.

Power spectral density (PSD) was estimated using a periodogram with a Hann window and density normalization. PSD was calculated separately for the left and right hemispheres. The bilateral measure was obtained by averaging the resulting spectra; hemispheric time series were not averaged before PSD estimation.

Absolute spectral power within a specified frequency band was defined as the integral of the PSD computed using the trapezoidal rule:

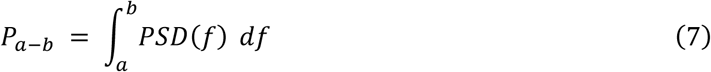

Reference power for normalization:

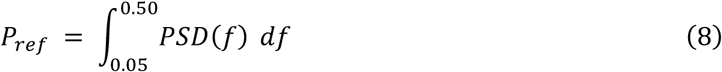

Relative power:

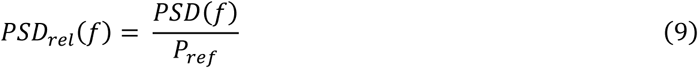

Relative band power for any selected frequency interval:

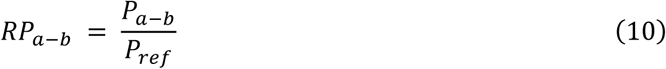

Expressed as a percentage:

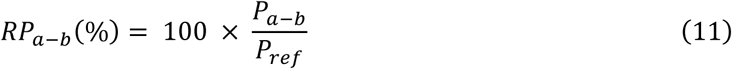

The primary spectral analysis was performed on 60-s REST segments. Given the segment duration, the lower boundary of quantitative analysis was set at 0.05 Hz, corresponding to three complete cycles within 60 s. Frequencies above 0.5 Hz were not analyzed because of the limited temporal bandwidth of the hemodynamic signal, reduced correspondence with neuronal activity above approximately 0.4 Hz, and the increasing contribution of physiological fluctuations, particularly those associated with respiration and cardiac activity (Ma et al., 2016b; Wright et al., 2017; Turley et al., 2017).

### Assessment of Lesion Area from WFOI Data

The cortical lesion was manually delineated at 10 min, 1.5 h, 1 day, and 3 days after the intervention. The recording acquired on the day of photothrombosis induction, before injury induction, was used as the individual baseline reference. Lesion boundaries were determined by jointly inspecting light-scattering images at 505 and 656 nm, pre-versus postinjury intensity-difference maps, maps of the temporal variability of HbT, and maps of homotopic functional connectivity. The final lesion mask was manually delineated using an interactive napari interface. Lesion area was calculated from the final binary mask using a spatial calibration of 22 µm/pixel in the full-resolution image space.

### Analysis of Spreading Depolarizations

Spreading depolarizations (SDs) were analyzed from 12-min WFOI recordings beginning at a median of 3.5 min (range, 3.3–5.5 min) after the end of laser irradiation. The morphology of the associated hemodynamic response was analyzed using HbT signals only for fully recorded SDs. For each event, response onset and offset and the HbT minimum corresponding to the initial vasoconstrictive phase were annotated. Events censored by recording boundaries or the onset of a subsequent SD were excluded from duration analysis. For construction of averaged HbT profiles, signals were smoothed with a P-spline and aligned in time to the HbT minimum and in amplitude to the signal level at event onset. Group curves were calculated after averaging multiple events within each animal. When several SDs occurred in one mouse, event parameters were first averaged within the animal, and these mouse-level values were then used for between-group analysis.

### Statistical Analysis

Statistical analyses were performed using custom Python scripts and the NumPy, pandas, SciPy, statsmodels, and Matplotlib libraries. The individual animal was the statistical unit; technical replicates, recordings, and segments were aggregated at the animal level before analysis. Missing values were not imputed, and statistical outlier exclusion was not performed. Unless otherwise stated, data are presented as mean ± standard deviation (std). Two-sided tests were used for t-tests and correlation analyses. A p-value < 0.05 was considered statistically significant; 95% confidence intervals (CIs) and effect sizes were also considered.

Two independent groups were compared using Welch’s t-test, and three groups were compared using one-way Welch’s analysis of variance (ANOVA). These tests were used as the primary methods for quantitative outcomes, given the small group sizes and without assuming equal variances. For pairwise comparisons, the difference in means, 95% CI, and Hedges’ g standardized effect size corrected for small-sample bias were reported. For descriptive interpretation, |g| < 0.2 was considered very small, 0.2–0.49 small, 0.5–0.79 moderate, and ≥0.8 large (Cohen, 1988).

Longitudinal outcomes were analyzed as within-animal changes: day 1 − baseline, day 3 − day 1, and day 3 − baseline. For WFOI-derived lesion area, day 1 − 1.5 h was also calculated. Within-group changes were compared with zero using one-sample t-tests. Holm’s correction was applied to families of multiple comparisons. Adjusted p-values were denoted p_adj_3 and p_adj_4, where the subscript indicates the number of tests in the family (three or four, respectively). Adjustments were applied, among other cases, to three pairwise between-group comparisons, tests of changes from zero in three groups, and regional comparisons across four cortical ROIs. Where adjustment was applied, statistical significance was determined from adjusted p-values.

Functional connectivity was analyzed statistically after Fisher z transformation, whereas Pearson correlation coefficients (r) were used to present the results. The number of spreading depolarizations was analyzed by Poisson regression; areas of capillary perfusion impairment and extravasation were compared using Welch’s t-test. Associations between outcomes were assessed using Spearman’s rank correlation coefficient (ρ), with 95% CIs obtained by percentile bootstrapping at the animal level (10,000 bootstrap resamples). For descriptive interpretation, |ρ| < 0.1 was regarded as negligible, 0.1–0.39 weak, 0.4–0.69 moderate, 0.7–0.89 strong, and ≥0.9 very strong (Schober et al., 2018). Correlation analyses were exploratory, and their p-values were not adjusted for multiple comparisons. Correlations were interpreted as associations rather than evidence of causality.

Absolute differences between percentages are expressed in percentage points (pp).

### Data Presentation in Figures

Quantitative plots show individual animal values, medians and interquartile ranges (IQRs) (boxplots), means (white diamonds), and mean ± std (black intervals), unless otherwise specified. Boxplot whiskers extend to the most extreme observations within 1.5 × IQR. In longitudinal plots, repeated measurements from the same animal are connected by lines. Functional connectivity maps are displayed on the Pearson r or Δr scale, whereas statistical analyses are performed in Fisher z space. Pairwise between-group comparisons (Welch’s t-tests with Holm’s correction) are indicated by brackets when the overall Welch’s ANOVA yields p < 0.055, unless otherwise specified. For visualization purposes, p-values < 0.055 are highlighted in red, whereas statistical significance is defined as p < 0.05.

### Use of Artificial Intelligence

ChatGPT (OpenAI, GPT-6 series) was used as an auxiliary tool for editing and debugging data-analysis code, language editing of the manuscript, transfer and formatting of numerical results from prepared statistical tables, and creation of selected illustrative elements of the experimental-design diagram. All numerical values, statistical results, code modifications, text, and graphic materials were checked and validated by the authors. Scientific conclusions and interpretation of the results were developed by the authors.

## Results

### Isoflurane Anesthesia Enhances Slow Hemodynamic Oscillations in the Sensorimotor Cortex

Visual inspection of videos showing Δ[HbT] changes revealed differences in the spatiotemporal organization of spontaneous hemodynamic activity. During wakefulness, diverse spatial patterns of signal fluctuations were observed, whereas under isoflurane anesthesia, recurrent episodes of widespread bilateral changes were recorded, predominantly in the somatosensory and primary motor cortices (Supplementary Video S1).

To quantify these differences, we analyzed the spectral power of spontaneous HbT changes in the 0.05–0.50 Hz range (Figure 2).

**Figure 2.**
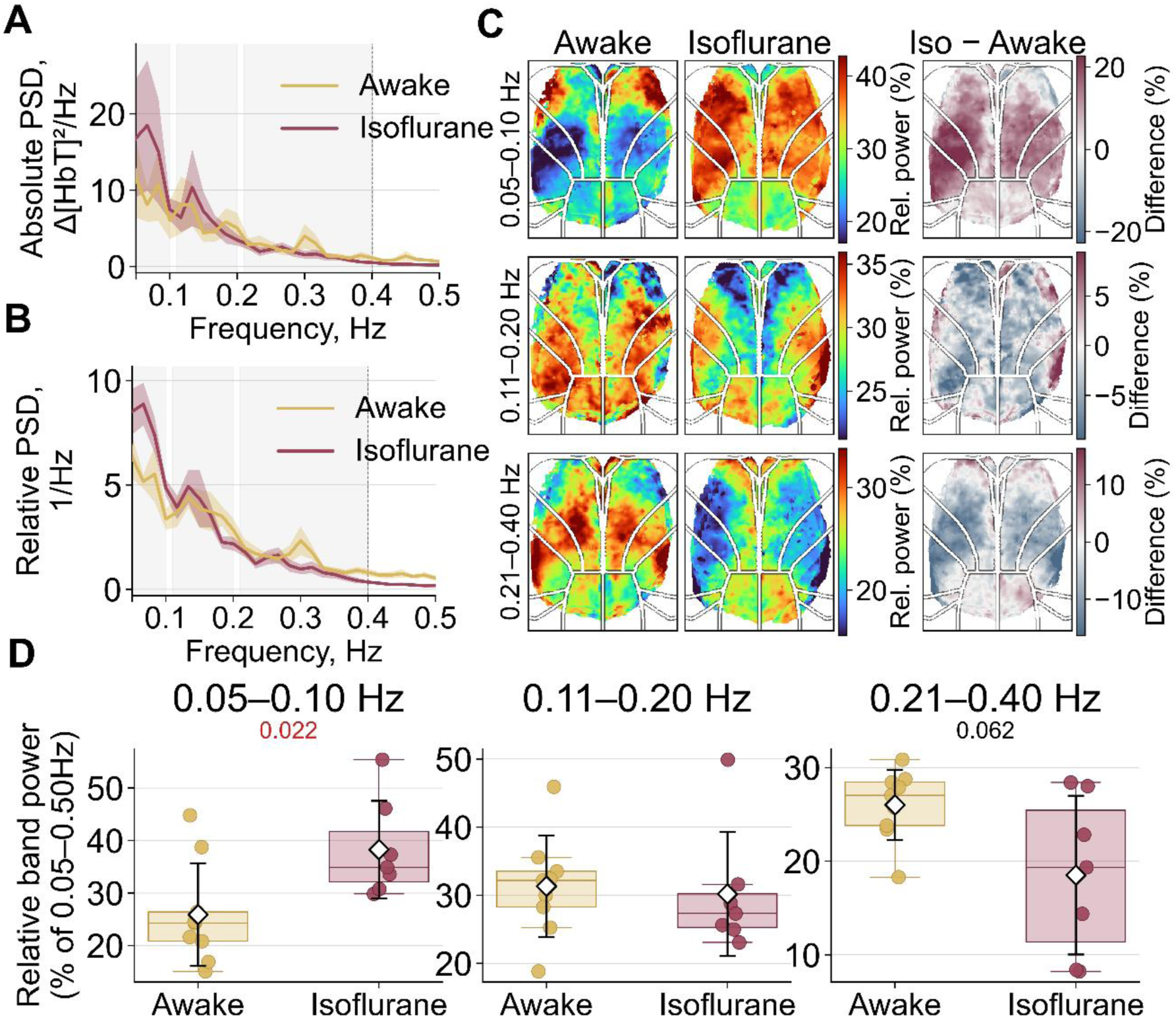
Isoflurane anesthesia alters the spectral characteristics of spontaneous hemodynamic activity, with a predominant increase in relative power at 0.05–0.10 Hz. (A,B) Absolute (A) and relative (B) power spectral density (PSD) of total hemoglobin concentration changes, Δ[HbT], averaged over the analyzed cortex during Awake (n = 9) and Isoflurane (n = 7) conditions. Values are shown as mean ± standard error of the mean (sem); shaded gray regions indicate the 0.05– 0.10, 0.11–0.20, and 0.21–0.40 Hz frequency bands. (C) Group-averaged pixelwise maps of relative spectral power within these bands, together with difference maps (Isoflurane − Awake, in percentage points). White outlines indicate boundaries of the analyzed cortical regions. (D) Relative spectral power in each band, averaged across the analyzed cortex and expressed as a percentage of total power in the 0.05–0.50 Hz range. Individual animal values, medians and interquartile ranges (boxplots), means (white diamonds), and std (black intervals) are shown. p-values were obtained using two-sided Welch’s t-tests.

Analysis of the absolute and relative power spectra of spontaneous Δ[HbT] fluctuations, averaged across the entire analyzed cortex, identified three frequency bands containing local spectral power peaks: 0.05–0.10, 0.11–0.20, and 0.21–0.40 Hz. Relative spectral power was then compared across these bands. The largest difference between conditions occurred at 0.05–0.10 Hz (Figure 2D): relative power was significantly higher in the Isoflurane condition (38.27 ± 9.28%; n = 7) than in the Awake condition (25.89 ± 9.81%; n = 9; Welch’s t-test, p = 0.022; g = 1.22). The increase in relative power at 0.05–0.10 Hz was accompanied by a numerical decrease at 0.21–0.40 Hz (p = 0.062), whereas differences in the 0.11–0.20 Hz band were less pronounced (Supplementary Table S1).

To characterize the spatial distribution of these frequency components, pixelwise maps of relative spectral power were constructed (Figure 2C). Under isoflurane anesthesia, increased 0.05–0.10 Hz power was observed predominantly in the somatosensory and primary motor cortices, where recurrent episodes of widespread bilateral signal changes had been noted during visual inspection. Regional analyses with Holm’s correction across four regions confirmed between-condition differences in SS and MOp (Supplementary Table S1).

### Isoflurane Anesthesia Increases MOp Homotopic Connectivity and Alters Functional Interactions between RSP and Other Cortical Regions

To evaluate cortical functional organization, we analyzed homotopic connectivity (HC), reflecting the coherence of hemodynamic fluctuations between symmetrically located regions of the two hemispheres, and global brain connectivity (GBC), representing the mean correlation between each cortical location and the remainder of the analyzed cortex. Both measures were derived from spontaneous Δ[HbT] changes after low-pass filtering with a cutoff frequency of 0.40 Hz (Figure 3).

**Figure 3.**
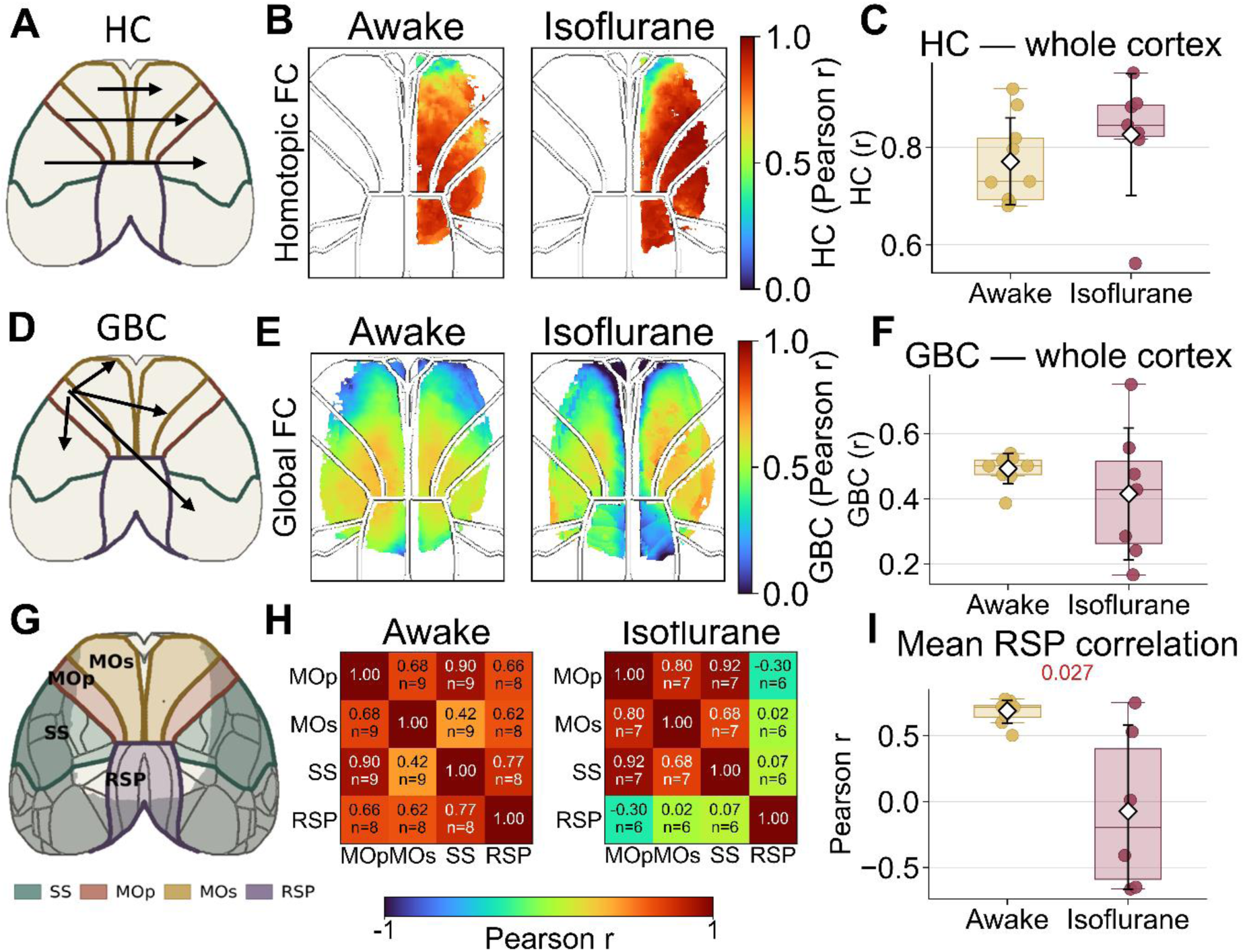
Isoflurane anesthesia increases homotopic connectivity in the primary motor cortex and alters functional interactions between the retrosplenial cortex and other regions. (A) Schematic of homotopic connectivity (HC) calculation. (B) Group-averaged HC maps during Awake (n = 9) and Isoflurane (n = 7) conditions. (C) Mean HC across the entire analyzed cortex. (D) Schematic of global brain connectivity (GBC) calculation. (E) Group-averaged GBC maps for Awake and Isoflurane conditions. (F) Mean GBC across the entire analyzed cortex. (G) Locations of analyzed cortical regions: primary motor cortex (MOp), secondary motor cortex (MOs), somatosensory cortex (SS), and retrosplenial cortex (RSP). (H) Correlation matrices of spontaneous hemodynamic fluctuations among these regions, averaged across animals. The sample size for each region pair is indicated within the corresponding matrix cell. (I) Mean correlation between RSP and MOp, MOs, and SS in Awake (n = 8) and Isoflurane (n = 6) conditions. Maps, plots, and matrices show Pearson r. In panel I, means (white diamonds) and mean ± std bounds (black intervals) were calculated in Fisher z space and back-transformed to Pearson r. The p-value was obtained using a two-sided Welch’s t-test in Fisher z space. FC, functional connectivity.

Mean HC across the entire analyzed cortex was high in both conditions and did not differ significantly: Pearson r was 0.771 ± 0.089 in Awake mice (n = 9) and 0.826 ± 0.125 under Isoflurane (n = 7; Welch’s t-test, p = 0.253; g = 0.60). No significant difference was found for GBC either: mean GBC Pearson r was 0.493 ± 0.046 in Awake mice (n = 9) and 0.415 ± 0.203 under Isoflurane (n = 7; Welch’s t-test, p = 0.523; g = −0.36). Statistical comparisons for both measures were performed in Fisher z space.

Regional analysis revealed significantly higher HC in MOp under isoflurane anesthesia: Pearson r was 0.923 ± 0.055 (n = 7) versus 0.786 ± 0.093 during wakefulness (n = 9). The Isoflurane − Awake difference in Fisher z space was 0.601 (95% CI, 0.265–0.937; Welch’s t-test, p_adj_4 = 0.0095; g = 1.95). Holm’s correction across four cortical regions was applied to regional comparisons. Full results are provided in Supplementary Table S1.

Visual inspection of Δ[HbT] recordings under isoflurane anesthesia also revealed episodes in which signal changes in the retrosplenial cortex (RSP) were opposite in direction to those in other analyzed regions. To quantify these interactions, correlation matrices were constructed and the mean correlation across three region pairs (MOs–RSP, MOp–RSP, and SS–RSP) was calculated in Fisher z space. Under Isoflurane, the mean correlation was negative (r = −0.072 (−0.667, 0.579); n = 6), whereas it was positive during wakefulness (r = 0.690 (0.594, 0.766); n = 8). Values represent back-transformed Fisher z means and mean ± std bounds. The Awake − Isoflurane difference in Fisher z space was 0.919 (95% CI, 0.151–1.687; Welch’s t-test, p = 0.027; g = 1.76). Thus, isoflurane anesthesia was associated with reduced concordance of spontaneous RSP hemodynamic fluctuations with those in the other cortical regions examined. This difference was also apparent in the group-level correlation matrices obtained by averaging individual animal matrices (Figure 3H).

### Induction of Isoflurane Anesthesia in Oxygen Increases Cortical Blood Oxygenation

To assess hemodynamic changes during induction of inhalational isoflurane anesthesia with oxygen as the carrier gas, we recorded the time courses of oxygenated (HbO), deoxygenated (HHb), and total (HbT) hemoglobin concentrations in the cerebral cortex in a separate experiment (Figure 4). For each animal, the difference between the mean values over the final 60 s and the first 7 s of continuous recording was calculated.

**Figure 4.**
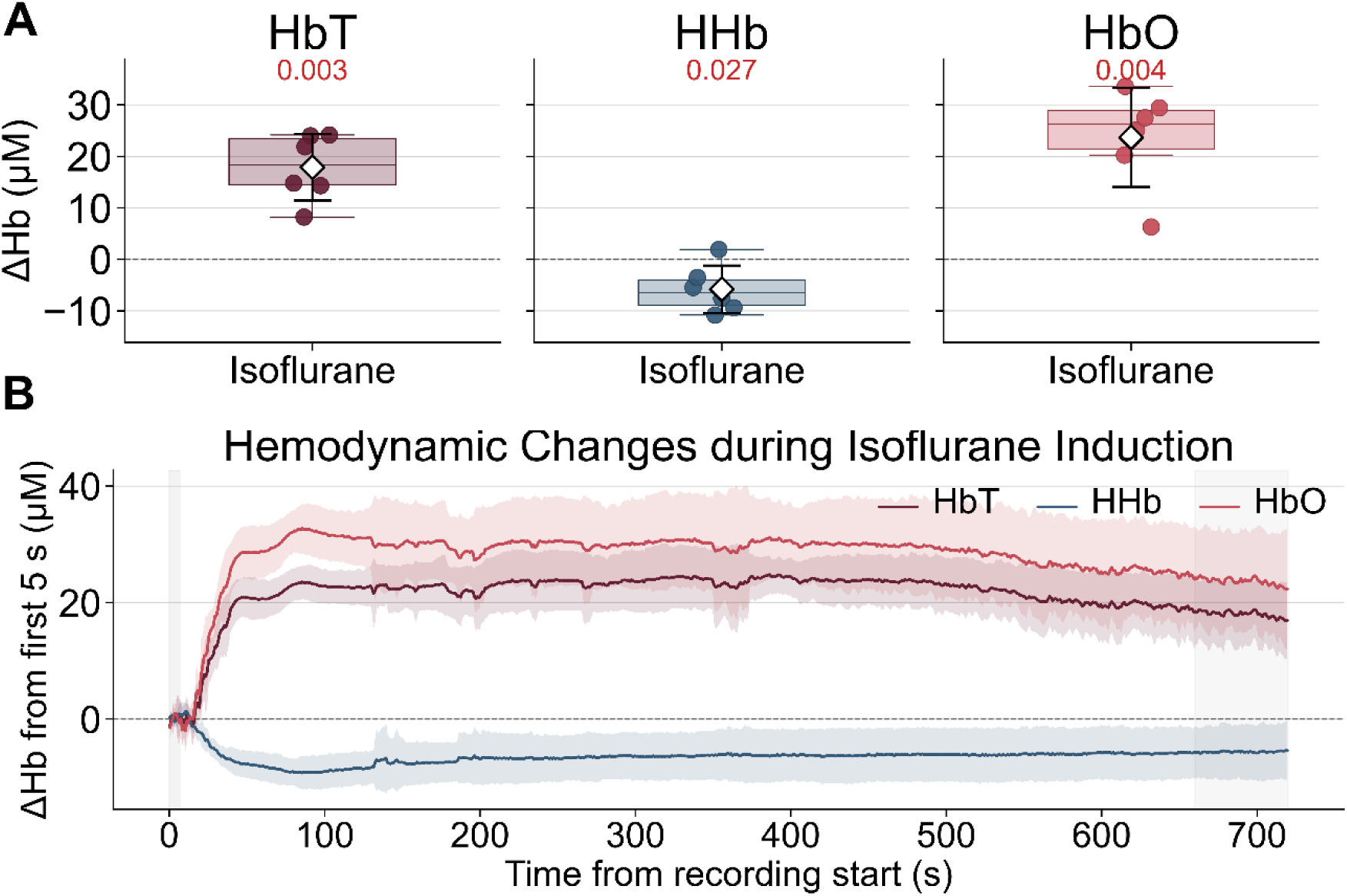
Changes in cortical hemoglobin concentrations during induction of inhalational isoflurane anesthesia in oxygen. (A) Differences in total (HbT), deoxygenated (HHb), and oxygenated (HbO) hemoglobin concentrations between the final 60 s and the first 7 s of recording. Individual animal values (n = 6) and boxplots are shown. p-values were obtained using one-sample t-tests with Holm’s correction across the three hemoglobin measures. (B) Time courses of HbT, HHb, and HbO changes averaged over the analyzed cortical surface. Curves represent mean values across six animals, and shaded areas indicate mean ± std. Changes are expressed relative to the mean signal during the first 5 s of recording. Time is measured from recording onset.

During anesthesia induction, HbO increased by 23.69 ± 9.61 µM (n = 6; 95% CI, 13.61–33.78; one-sample t-test, p_adj_3 = 0.004), accompanied by a decrease in HHb of −5.79 ± 4.60 µM (95% CI, −10.62 to −0.96; p_adj_3 = 0.027). HbT simultaneously increased by 17.90 ± 6.46 µM (p_adj_3 = 0.003; Figure 4A). Holm’s correction was applied across the three hemoglobin measures.

Analysis of the temporal dynamics showed that the increase in HbO and HbT and the decrease in HHb occurred primarily at the beginning of the recording (Figure 4B). These changes persisted thereafter, although HbO and HbT gradually declined from their peak values toward the end of the recording.

Thus, induction of inhalational isoflurane anesthesia in oxygen was accompanied by increased cortical blood oxygenation and elevated HbT, with both changes persisting throughout the recording.

### Assessment of Changes Induced by Focal Cortical Injury

We next assessed the severity and functional consequences of focal cortical injury under three experimental conditions: laser exposure without effective intravenous administration of the photosensitizer (Laser), photothrombosis in awake animals (Awake PT), and photothrombosis under isoflurane anesthesia (IsO_2_ PT; 1.5% isoflurane with oxygen as the carrier gas). Longitudinal measurements from 1.5 h after injury onward were performed in awake animals, enabling assessment of the consequences of injury-induction conditions without the direct effects of isoflurane during recording.

### Differences in Cortical Lesion Area Are Detectable within 10 Minutes of Injury Induction

The severity of focal cortical injury was evaluated using two complementary measures of lesion area: WFOI at 10 min, 1.5 h, day 1, and day 3 after injury, and TTC staining of brain tissue on day 3. In WFOI data, the injured region primarily encompassed the somatosensory cortex (SS) and part of the primary motor cortex (MOp); with more severe injury, it extended mainly toward MOp and, to a lesser degree, into MOs and RSP (Figure 5A).

**Figure 5.**
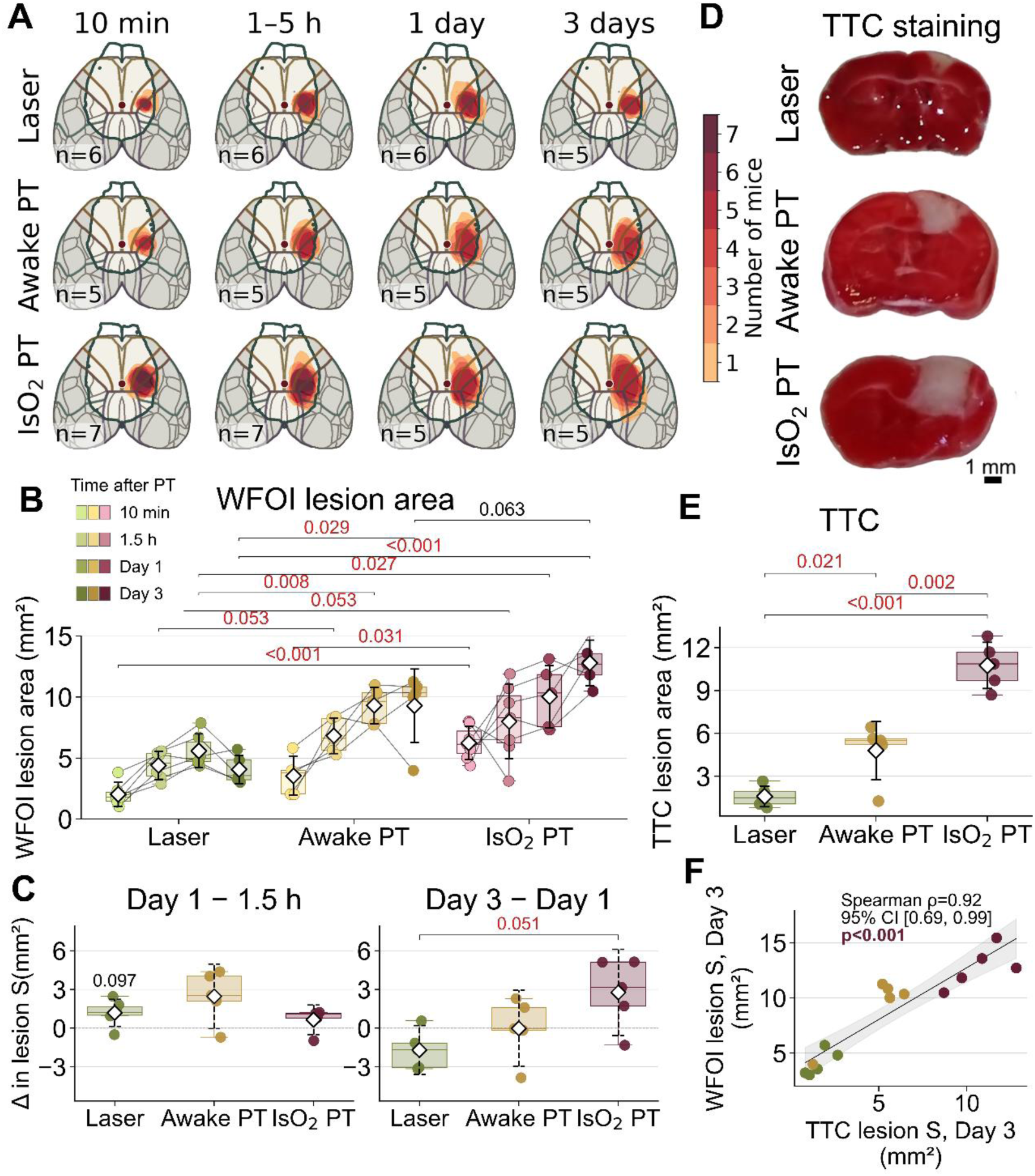
Cortical lesion size varies across experimental conditions and shows concordance between WFOI and TTC assessments. (A) WFOI-derived lesion maps in the Allen Mouse Brain Atlas coordinate space; the outline indicates the cortical area available for analysis, and the color scale indicates the number of animals with injury at each location. (B) Individual trajectories of lesion area at 10 min, 1.5 h, day 1, and day 3 after irradiation in Laser, Awake PT, and IsO₂ PT (n = 5 per group for complete longitudinal trajectories). (C) Changes in lesion area between 1.5 h and day 1 and between days 1 and 3. (D) Representative TTC-stained coronal sections. Scale bar, 1 mm. (E) TTC-negative lesion area on day 3 (n = 5 per group). (F) Association between WFOI-and TTC-derived lesion areas on day 3 (n = 15), showing Spearman’s ρ, bootstrap 95% CI, and p-value. Numeric labels above boxplots in panel C indicate Holm-adjusted p-values from one-sample t-tests comparing changes with zero. Brackets denote pairwise Welch’s t-tests with Holm’s correction. Black dashed intervals in panel C indicate 95% CIs of the mean.

Between-group differences in lesion area were apparent as early as 10 min after irradiation (Welch’s ANOVA, p < 0.001). Lesion areas were 2.03 ± 0.99 mm² in Laser (n = 6), 3.54 ± 1.58 mm² in Awake PT (n = 5), and 6.24 ± 1.36 mm² in IsO_2_ PT (n = 7) (Figure 5B). Lesion area in IsO_2_ PT was larger than in Awake PT (difference, 2.70 mm²; 95% CI, 0.68–4.72; Welch’s t-test, p_adj_3 = 0.031; g = 1.71) and Laser (difference, 4.22 mm²; 95% CI, 2.77–5.66; p_adj_3 < 0.001; g = 3.26). The mean lesion area was also larger in Awake PT than in Laser, with a large standardized effect size (p_adj_3 = 0.109; g = 1.08). Holm’s correction was applied to the three pairwise group comparisons separately at each time point (here and below, unless otherwise specified).

At 1.5 h, lesion areas were 4.41 ± 1.16 mm² in Laser (n = 6), 6.84 ± 1.46 mm² in Awake PT (n = 5), and 8.00 ± 3.03 mm² in IsO_2_ PT (n = 7). The overall between-group difference persisted (Welch’s ANOVA, p = 0.016); with larger mean lesion areas in both photothrombosis groups than in Laser (Welch’s t-test, all p_adj_3 ≥ 0.053).

By day 1, lesion areas were 5.60 ± 1.38 mm² in Laser (n = 6), 9.31 ± 1.49 mm² in Awake PT (n = 5), and 10.04 ± 2.57 mm² in IsO_2_ PT (n = 5; Welch’s ANOVA, p = 0.005). Both photothrombosis groups differed from Laser (Awake PT: p_adj_3 = 0.008, g = 2.37; IsO_2_ PT: p_adj_3 = 0.027, g = 2.03), but not from each other (p_adj_3 = 0.602; g = 0.31). Changes in lesion area between 1.5 h and day 1 were +1.19 ± 0.99, +2.47 ± 2.02, and +0.66 ± 0.92 mm², respectively (n = 6, 5, and 5); these changes did not differ significantly among groups (Welch’s ANOVA, p = 0.257).

The trajectories diverged most strongly between days 1 and 3: lesion area decreased in Laser (−1.69 ± 1.53 mm²), remained nearly unchanged in Awake PT (−0.02 ± 2.38 mm²), and increased in IsO_2_ PT (+2.76 ± 2.69 mm²; n = 5 per group). The overall comparison of changes revealed between-group differences (Welch’s ANOVA, p = 0.045). The largest difference occurred between IsO_2_ PT and Laser (4.45 mm²; Welch’s t-test, p_adj_3 = 0.051; g = 1.84).

By day 3, lesion areas were 4.07 ± 1.16 mm² in Laser, 9.29 ± 3.00 mm² in Awake PT, and 12.80 ± 1.87 mm² in IsO_2_ PT (n = 5 per group; Welch’s ANOVA, p < 0.001). Both photothrombosis groups had significantly larger lesions than Laser (Awake PT: p_adj_3 = 0.029, g = 2.07; IsO2 PT: p_adj_3 < 0.001, g = 5.08). The mean lesion area was also larger in IsO_2_ PT than in Awake PT, with a large standardized effect size (p_adj_3 = 0.063; g = 1.27). Thus, by day 3, mean lesion areas showed a graded increase: Laser < Awake PT < IsO_2_ PT. Full results for absolute lesion area and its longitudinal changes are provided in Supplementary Tables S5A–S5D.

### TTC Staining Confirms Differences in Injury Severity among Groups on Day 3

TTC staining of coronal sections at the end of the experiment provided an independent assessment of the extent of injury through the depth of the tissue (Figure 5D,E). In representative sections, the lesion extended into the cortical depth in all groups. In Laser, the TTC-negative area was more restricted and narrowed with depth, producing a predominantly conical profile, whereas photothrombosis involved a wider region of tissue.

On day 3, the TTC-negative lesion area was 1.60 ± 0.73 mm² in Laser, 4.81 ± 2.03 mm² in Awake PT, and 10.75 ± 1.61 mm² in IsO_2_ PT (n = 5 per group; Welch’s ANOVA, p < 0.001; Figure 5E). Pairwise Welch’s t-tests with Holm’s correction across the three group comparisons showed that the area in Awake PT exceeded that in Laser by 3.21 mm² (p_adj_3 = 0.021; g = 1.90), whereas that in IsO_2_ PT exceeded Laser by 9.15 mm² (p_adj_3 < 0.001; g = 6.60). Moreover, the TTC-negative area in IsO_2_ PT was larger than in Awake PT by 5.94 mm² (95% CI, 3.24–8.64; p_adj_3 = 0.002; g = 2.93). Thus, TTC staining confirmed differences in structural injury severity among all three groups.

Despite the different measurement principles, the WFOI-derived surface lesion area on day 3 was positively correlated with the TTC-negative lesion area measured in a coronal section (Spearman’s ρ = 0.921; n = 15; 95% bootstrap CI, 0.693–0.985; p < 0.001; Figure 5F). To compare the two methods more directly, maximum lesion diameters determined by WFOI and TTC staining on day 3 were also compared. These measures likewise showed a strong positive correlation (ρ = 0.788; n = 15; 95% bootstrap CI, 0.452–0.945; p = 0.000482). These findings indicate concordance between the methods in estimating relative injury severity, although lesion areas were measured in different anatomical planes.

Together, both methods showed that differences among groups were evident both at the cortical surface and through the depth of the tissue, with the most extensive injury following photothrombosis induced under isoflurane anesthesia with oxygen as the carrier gas. Laser irradiation without effective intravenous photosensitizer delivery also produced a lesion, although of smaller size. Overall, injury severity increased in the order Laser < Awake PT < IsO_2_ PT.

### Homotopic Functional Connectivity Decreases within 1.5 Hours of Injury, Most Markedly in IsO2 PT

Homotopic connectivity (HC) reflects interactions between symmetrical regions of the two cerebral hemispheres, partly supported by callosal connections, and is disrupted by focal ischemic brain injury (Bauer et al., 2014). To assess postinjury changes, we analyzed HC based on spontaneous Δ[HbT] changes low-pass filtered at 0.40 Hz, both across the available cortex and separately in MOp, MOs, SS, and RSP (Figure 6).

**Figure 6.**
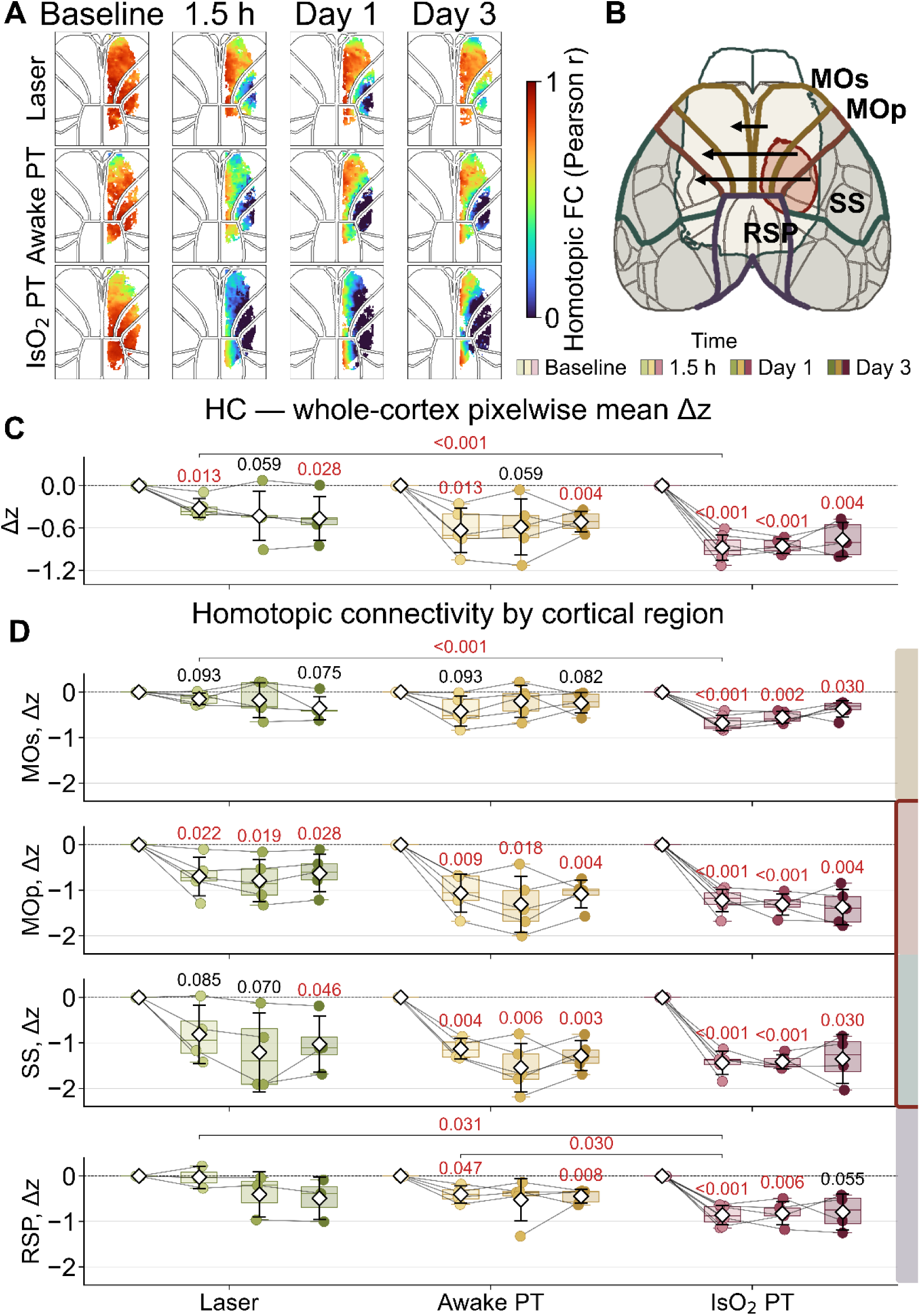
Focal cortical injury is accompanied by a predominant decrease in homotopic connectivity in SS and MOp. (A) Group-averaged HC maps before injury and at 1.5 h, day 1, and day 3 afterward for Laser, Awake PT, and IsO_2_ PT. Maps show Pearson r. (B) Schematic of pixelwise HC calculation. (C) Change in mean HC across the analyzed cortex relative to baseline (Fisher Δz). (D) Changes in HC in MOp, MOs, SS, and RSP. Actual sample sizes ranged from 3 to 7 animals, depending on the region and time point. Numeric labels above boxplots indicate Holm-adjusted p-values from one-sample t-tests comparing Δ with zero. Brackets indicate pairwise Welch’s t-tests with Holm’s correction across three group comparisons (C) or four cortical regions (D). Thus, HC detected functional changes within 1.5 h after injury, including in the group with the mildest lesions. Regional analysis also revealed differences among injury-induction conditions not only near the principal lesion but also in less affected cortical regions, particularly MOs and RSP.

Before injury, mean whole-cortex HC (Pearson r) was 0.814 ± 0.068 in Laser (n = 6), 0.763 ± 0.059 in Awake PT (n = 5), and 0.769 ± 0.085 in IsO_2_ PT (n = 7). There were no statistically significant baseline differences among groups (Welch’s ANOVA, p = 0.454). Subsequent changes were expressed relative to each animal’s baseline in Fisher z space (Δz).

At 1.5 h after injury, mean whole-cortex HC had decreased in all three groups: Δz = −0.317 ± 0.135 in Laser (n = 5), −0.633 ± 0.311 in Awake PT (n = 5), and −0.881 ± 0.179 in IsO_2_ PT (n = 7) (Figure 6C). The decrease from baseline was statistically significant in all groups, including Laser, which had the smallest lesions (one-sample t-test, p_adj_3 = 0.013, 0.013, and <0.001, respectively). The magnitude of the early change also differed among groups (Welch’s ANOVA, p = 0.001). The decrease in IsO_2_ PT was greater than in Laser (difference, 0.563 Fisher Δz; 95% CI, 0.361–0.765; Welch’s t-test, p_adj_3 < 0.001; g = 3.20). The remaining pairwise differences were not statistically significant (p_adj_3 = 0.175).

By day 1, reduced HC relative to baseline persisted: Δz = −0.430 ± 0.348 in Laser, −0.584 ± 0.394 in Awake PT, and −0.857 ± 0.101 in IsO2 PT (n = 5 per group). The largest reduction was observed in IsO2 PT (one-sample t-test, p_adj_3 < 0.001), with the mean HC changes retaining the same graded pattern across groups (Welch’s ANOVA, p = 0.081).

No substantial recovery of mean HC was observed between days 1 and 3. By day 3, reductions from baseline persisted in all groups: Δz = −0.465 ± 0.308 in Laser, −0.509 ± 0.142 in Awake PT, and −0.764 ± 0.243 in IsO_2_ PT (n = 5 per group; one-sample t-test, p_adj_3 = 0.028, 0.004, and 0.004, respectively). The magnitude of the change did not differ significantly among groups (Welch’s ANOVA, p = 0.189).

Regional analysis revealed spatially heterogeneous HC changes (Figure 6D). The largest decreases occurred in SS and adjacent MOp, which corresponded to the main lesion region. However, between-group differences were also detected as early as 1.5 h in the less affected MOs and RSP. In MOs, HC changes were Δz = −0.146 ± 0.124 in Laser (n = 5), −0.421 ± 0.331 in Awake PT (n = 5), and −0.680 ± 0.167 in IsO_2_ PT (n = 7). The decrease in IsO2 PT was greater than in Laser (difference, 0.534 Fisher Δz; 95% CI, 0.346–0.722; Welch’s t-test, p_adj_4 < 0.001; g = 3.25). In RSP, the early HC change was Δz = −0.027 ± 0.244 in Laser (n = 3), −0.403 ± 0.188 in Awake PT (n = 4), and −0.860 ± 0.212 in IsO_2_ PT (n = 7). The decrease in IsO2 PT was greater than in Laser (Welch’s t-test, p_adj_4 = 0.031; g = 3.41) and Awake PT (p_adj_4 = 0.030; g = 2.04). At later time points, the largest HC reductions persisted in SS and MOp. By day 3, reductions from baseline were statistically significant in all three groups, including Laser (one-sample t-test, p_adj_3 < 0.05). Changes in MOs were less pronounced, whereas RSP showed between-group differences in the magnitude and persistence of the decrease. Full regional results are provided in Supplementary Table S2.

### Global Brain Connectivity in the Intact Hemisphere Is Less Affected by Cortical Injury than Homotopic Connectivity

Global brain connectivity (GBC) characterizes the similarity of spontaneous hemodynamic fluctuations at each cortical location to those in the remaining analyzed region. Here, GBC was calculated in the hemisphere contralateral to the lesion to assess functional interactions within morphologically intact cortex (Figure 7). Baseline values are reported as Pearson r, whereas subsequent changes were analyzed in Fisher z space (Δz).

**Figure 7.**
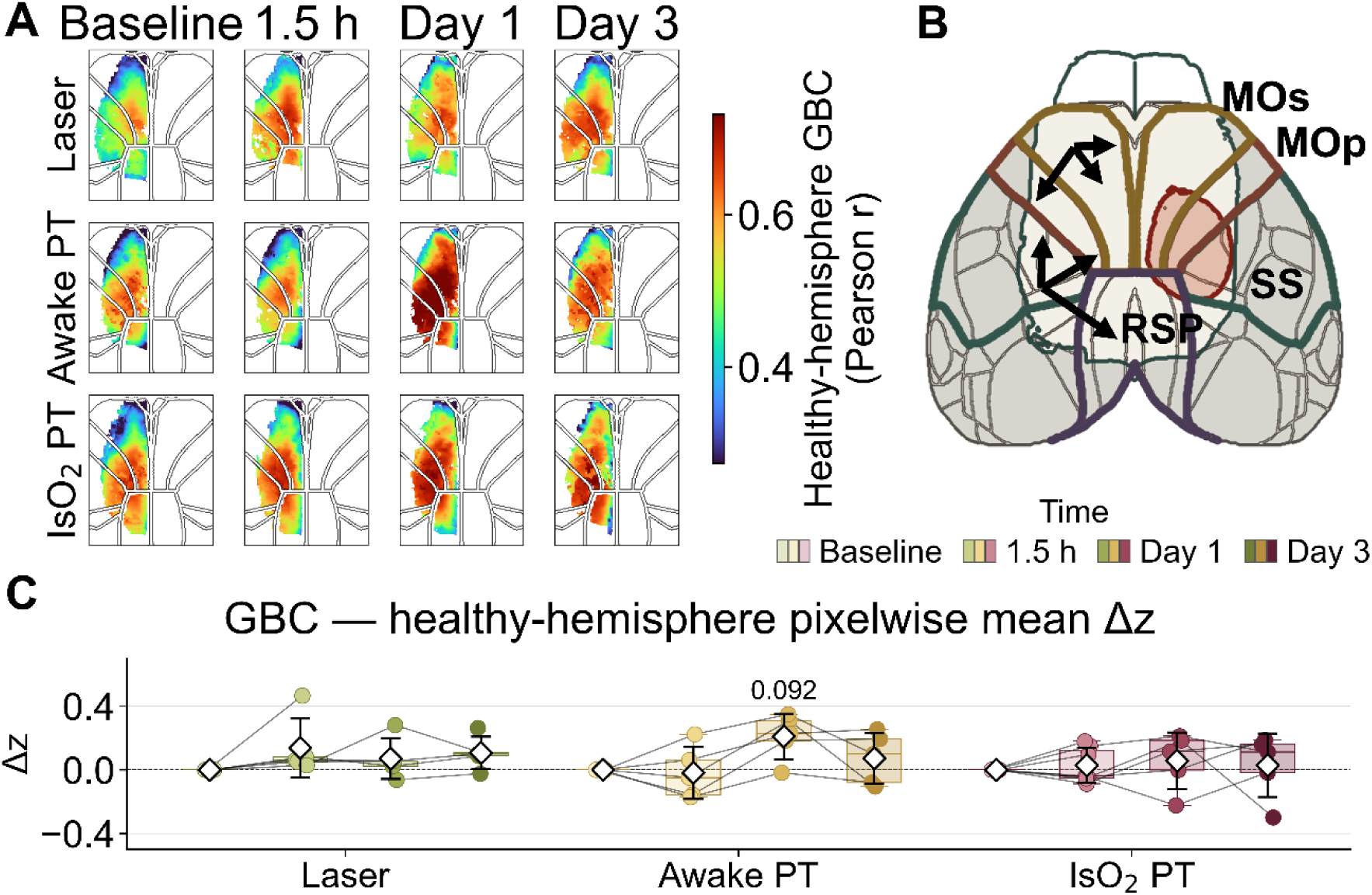
Focal cortical injury is associated with transient changes in global brain connectivity in the intact hemisphere, most evident in Awake PT. (A) Group-averaged GBC maps of the hemisphere contralateral to the lesion before injury and at 1.5 h, day 1, and day 3 afterward in Laser, Awake PT, and IsO_2_ PT. Maps show Pearson r. (B) Schematic of GBC calculation. (C) Change in mean GBC of the intact hemisphere relative to baseline (Fisher Δz). Sample sizes ranged from 5 to 7, depending on group and time point. Numeric labels above boxplots indicate Holm-adjusted p-values from one-sample t-tests comparing Δ with zero. Brackets indicate pairwise Welch’s t-tests with Holm’s correction across three group comparisons (C).

Before injury, mean GBC was 0.456 ± 0.070 in Laser (n = 6), 0.491 ± 0.135 in Awake PT (n = 5), and 0.512 ± 0.067 in IsO_2_ PT (n = 7; Welch’s ANOVA, p = 0.407). Postinjury changes in GBC were less pronounced than those in HC. The largest increase in mean GBC was observed in Awake PT (Δz = +0.210 ± 0.143; n = 5; one-sample t-test, p_adj_3 = 0.092) on day 1. By day 3, mean GBC values in all groups were closer to their respective baseline levels than the corresponding HC values. No statistically significant between-group differences in GBC changes were detected at any time point.

Regional analysis revealed the most pronounced day-1 changes in motor regions. In particular, GBC in MOs of the Awake PT group showed a numerical increase relative to baseline (Δz = +0.213 ± 0.120; n = 5; one-sample t-test, p_adj_3 = 0.051), which was no longer evident on day 3. Full results are provided in Supplementary Table S3.

Thus, in contrast to the persistent reduction in HC, GBC in the contralateral hemisphere showed comparatively modest changes, with a transient numerical increase most evident in Awake PT on day 1.

### Cortical Injury Causes Motor Deficits and Increases NSE, Most Prominently in IsO2 PT

As independent indicators of the consequences of cortical injury, we assessed motor deficits in the Grid and Cylinder tests and measured serum NSE and S100β concentrations (Figure 8; Supplementary Tables S4A and S4B).

**Figure 8.**
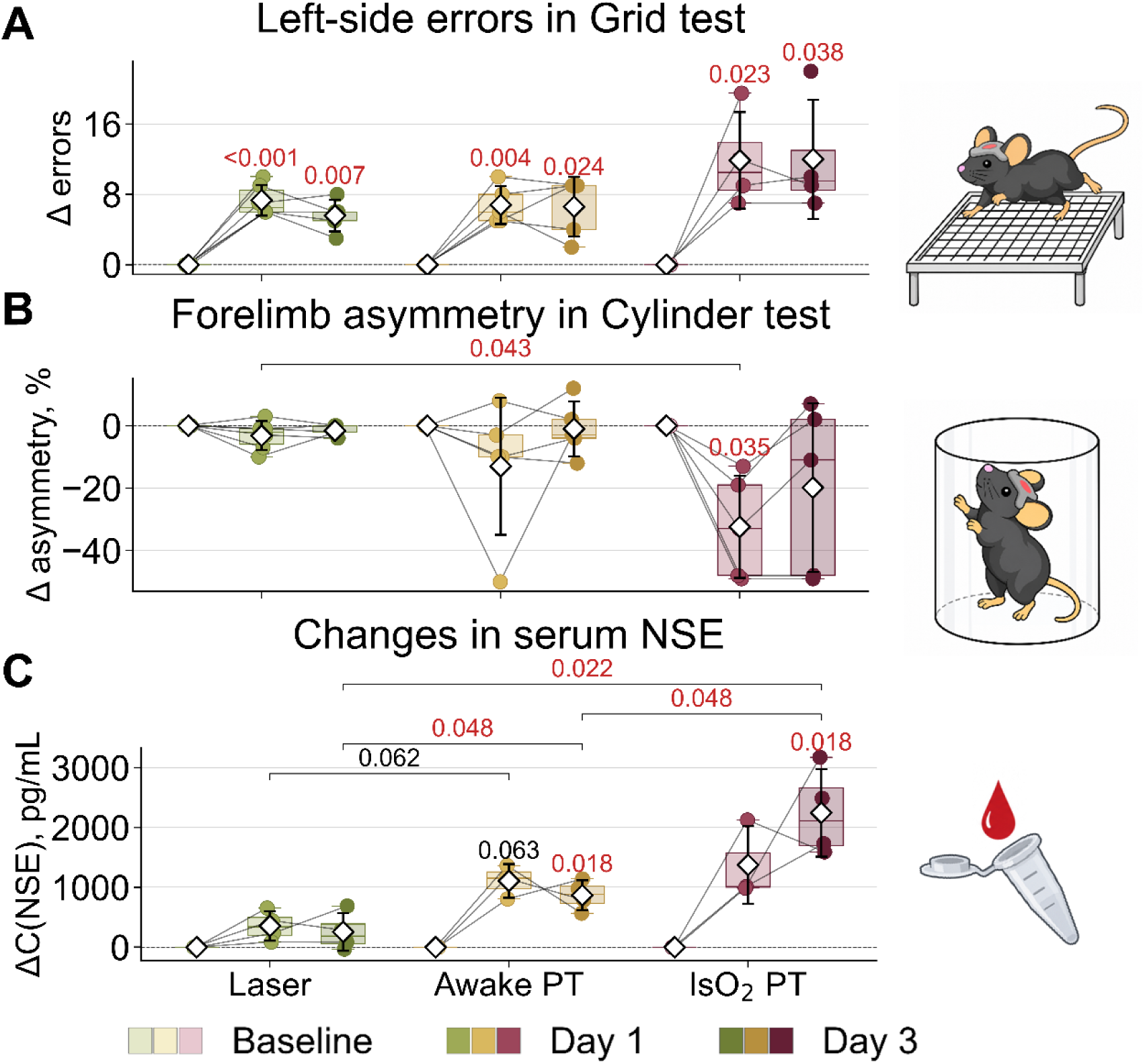
Motor and biochemical consequences of focal cortical injury. Individual changes relative to baseline (Δ = postinjury − preinjury) are shown. (A) Change in the number of left-limb errors in the Grid test. (B) Change in the asymmetry of forelimb use in the Cylinder test (pp). (C) Change in serum neuron-specific enolase (NSE) concentration at 1 and 3 days after injury. Actual sample sizes ranged from 3 to 6 depending on outcome and time point. Numeric labels above boxplots indicate Holm-adjusted p-values from one-sample t-tests comparing Δ with zero.

In the Grid test, no baseline between-group differences were detected (Welch’s ANOVA, p = 0.803; Figure 8A). By day 1, the number of left-limb errors had increased in all groups: by +7.3 ± 1.8 in Laser (n = 6; here and below, one-sample t-test, p_adj_3 < 0.001), +6.8 ± 2.2 in Awake PT (n = 5; p_adj_3 = 0.004), and +11.9 ± 5.5 in IsO_2_ PT (n = 4; p_adj_3 = 0.023). There was no significant between-group difference in the magnitude of change (Welch’s ANOVA, p = 0.323). By day 3, error counts remained elevated in all groups: +5.6 ± 1.8 (n = 5; p_adj_3 = 0.007), +6.6 ± 3.4 (n = 5; p_adj_3 = 0.024), and +12.0 ± 6.8 (n = 4; p_adj_3 = 0.038), respectively, with no significant group difference in the magnitude of change (Welch’s ANOVA, p = 0.287). Thus, the Grid test detected even relatively mild injury but discriminated poorly between levels of injury severity across groups.

In the Cylinder test, baseline values likewise did not differ among groups (Welch’s ANOVA, p = 0.118; Figure 8B). By day 1, the change in the asymmetry measure was −3.2 ± 4.7 pp in Laser (n = 6), −13.0 ± 22.0 pp in Awake PT (n = 5), and −32.4 ± 16.4 pp in IsO_2_ PT (n = 5). A significant change from baseline was detected in IsO2 PT (one-sample t-test, p_adj_3 = 0.035), and the magnitude of change differed among groups (Welch’s ANOVA, p = 0.028). The largest difference was between IsO_2_ PT and Laser (difference, −29.2 pp; 95% CI, −49.31 to −9.15; Welch’s t-test, p_adj_3 = 0.043; g = −2.33). Between days 1 and 3, values in the photothrombosis groups shifted toward baseline, but no within-group recovery reached statistical significance (one-sample t-test, all p_adj_3 > 0.05). The magnitude of recovery also did not differ among groups (Welch’s ANOVA, p = 0.174). Thus, the Cylinder test was less sensitive to mild laser-induced injury but detected differences in early motor impairment across injury-induction conditions.

Baseline NSE concentrations did not differ among groups (Welch’s ANOVA, p = 0.547; Figure 8C). On day 1, NSE changes were +355 ± 248 pg/mL in Laser (n = 4), +1107 ± 282 pg/mL in Awake PT (n = 3), and +1375 ± 649 pg/mL in IsO_2_ PT (n = 3; Welch’s ANOVA, p = 0.051). Compared with Laser, both photothrombosis groups showed substantially greater mean increases in NSE, with large standardized effect sizes (Awake PT: difference, +752 pg/mL; Welch’s t-test, p_adj_3 = 0.062; g = 2.42; IsO_2_ PT: difference, +1020 pg/mL; p_adj_3 = 0.201; g = 1.89).

By day 3, NSE changes were +256 ± 317 pg/mL in Laser, +869 ± 248 pg/mL in Awake PT, and +2245 ± 733 pg/mL in IsO_2_ PT (n = 4 per group). Increases relative to baseline were statistically significant in both photothrombosis groups (one-sample t-test, p_adj_3 = 0.018 for each), but not in Laser. The magnitude of change differed among groups (Welch’s ANOVA, p = 0.010). Compared with Laser, the increase was greater in Awake PT (difference, +613 pg/mL; 95% CI, 114–1112; Welch’s t-test, p_adj_3 = 0.048; g = 1.87) and IsO_2_ PT (+1989 pg/mL; 95% CI, 890–3089; p_adj_3 = 0.022; g = 3.06). Furthermore, the increase in IsO2 PT exceeded that in Awake PT (+1376 pg/mL; 95% CI, 264–2488; p_adj_3 = 0.048; g = 2.19).

Serum S100β concentration was also assessed (Supplementary Tables S4A and S4B). Unlike NSE, S100β showed no statistically confirmed increase after injury.

Thus, the outcomes assessed reflected different aspects of cortical injury. The Grid test detected persistent motor deficits even in Laser, whereas the Cylinder test revealed greater early impairment in IsO_2_ PT. By day 3, changes in NSE differentiated all three groups, reflecting the graded severity of cortical injury.

### Cortical Lesion Area Is Associated with Early HC Reduction in MOs and the Severity of Motor and Biochemical Impairments

To assess associations between functional consequences and structural injury severity, TTC-negative lesion area on day 3 was compared with the HC change in MOs at 1.5 h, residual increases in left-limb errors in the Grid test, and changes in NSE concentration by day 3 (Figure 9).

**Figure 9.**
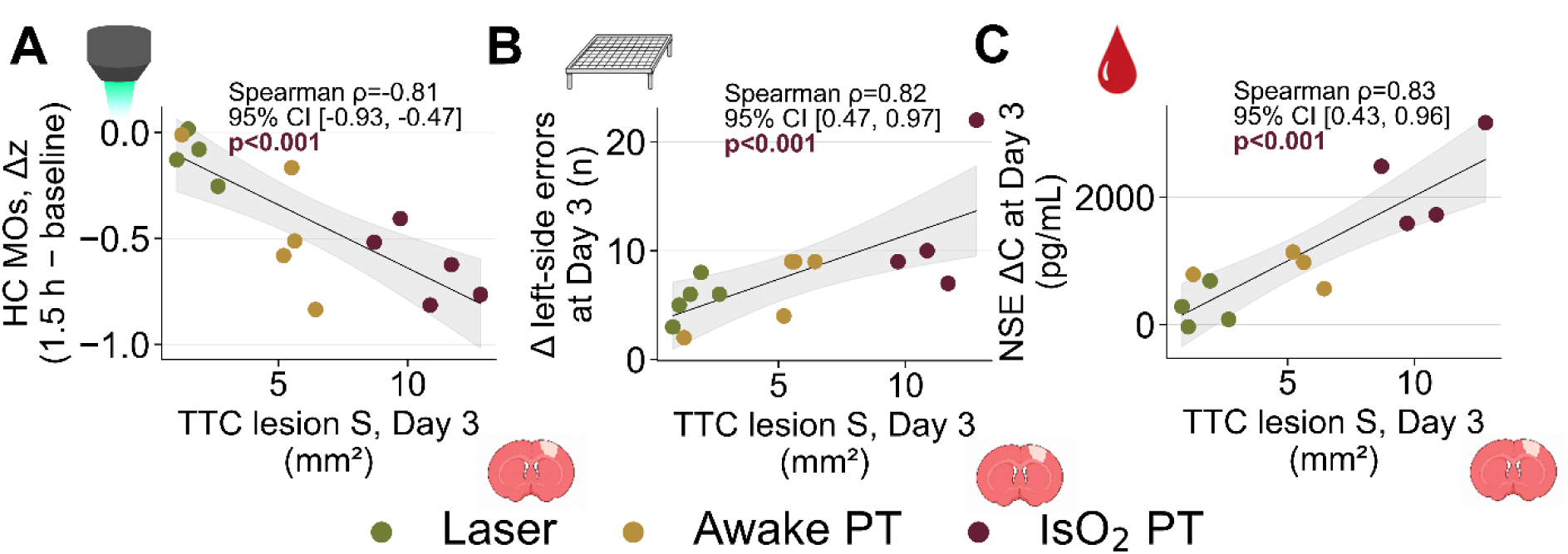
Associations between cortical lesion size and functional, motor, and biochemical changes. TTC-negative lesion area on day 3 was compared with changes from each animal’s baseline. (A) Homotopic connectivity (HC) in the secondary motor cortex (MOs) at 1.5 h (Fisher Δz; n = 14). (B) Left-limb error count in the Grid test on day 3 (n = 14). (C) Serum neuron-specific enolase (NSE) concentration on day 3 (n = 12). Spearman’s correlation coefficients (ρ), bootstrap 95% CIs, and unadjusted p-values are shown. Fitted linear regression lines and their 95% confidence bands visualize the direction of each association.

A strong negative correlation was observed between TTC-negative lesion area on day 3 and the early HC change in MOs (Spearman’s ρ = −0.811; n = 14; 95% bootstrap CI, −0.933 to −0.474; p < 0.001; Figure 9A). Animals with larger lesions on day 3 showed a greater decrease in HC from baseline as early as 1.5 h after injury.

Positive correlations of similar absolute magnitude were observed between TTC-negative lesion area and the residual increase in Grid-test errors (ρ = 0.817; n = 14; 95% bootstrap CI, 0.472–0.966; p < 0.001; Figure 9B) and the increase in NSE (ρ = 0.825; n = 12; 95% bootstrap CI, 0.435–0.957; p < 0.001; Figure 9C). Thus, greater structural injury was associated with more pronounced early HC reduction in MOs and more severe motor and biochemical consequences by day 3.

### Photothrombosis under Isoflurane Anesthesia with Oxygen Produces More Extensive Impairments in Capillary Perfusion and BBB Permeability

To further characterize vascular consequences of photothrombosis, the distribution of intravenously administered 10-kDa FITC-labeled dextran was assessed 1.5 h after injury. The area lacking capillary filling was determined from first-pass recordings, whereas the area of dextran extravasation, used as an indicator of BBB permeability disruption, was determined from images acquired 1 min after administration (Figure 10). Awake PT and IsO_2_ PT were compared; no comparable measurements after intravenous dextran administration were available for Laser.

**Figure 10.**
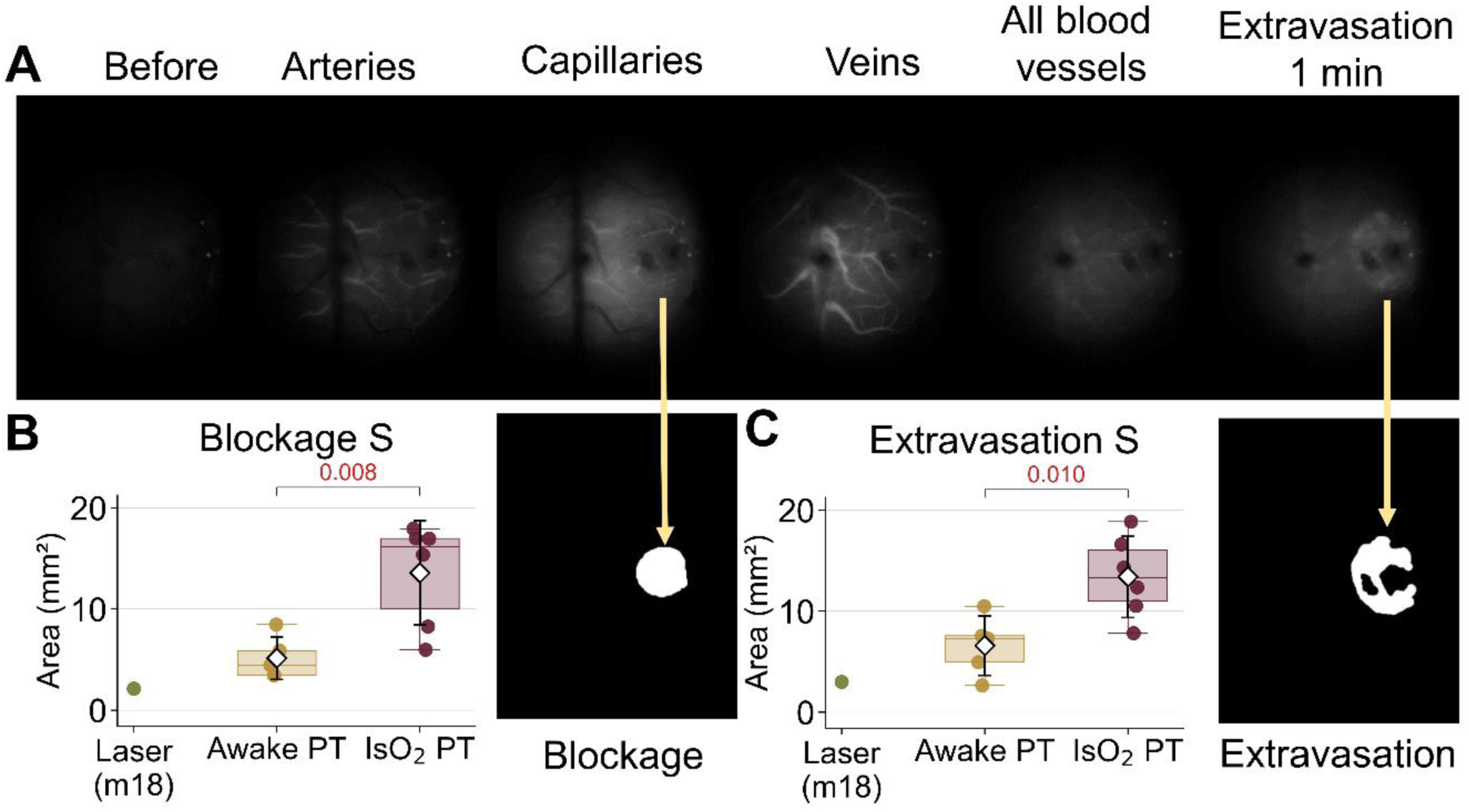
Photothrombosis under isoflurane anesthesia is associated with greater impairment of capillary perfusion and BBB permeability. (A) Representative sequence of the first pass of intravenously administered 10-kDa FITC-dextran through the cortical vasculature and extravasation at 1 min. (B) Area of impaired capillary perfusion in Awake PT (n = 5) and IsO_2_ PT (n = 6), with an example mask. (C) FITC-dextran extravasation area in the same groups, with an example mask. p-values were obtained using two-sided Welch’s t-tests. The Laser-group animal (m18) that received saline instead of Rose Bengal is shown only as an illustrative reference and was not included in statistical comparisons.

The area lacking capillary perfusion was 5.15 ± 2.11 mm² in Awake PT (n = 5) and 13.59 ± 5.13 mm² in IsO2 PT (n = 6; Figure 10B). This area was larger by 8.44 mm² in IsO_2_ PT (Welch’s t-test, p = 0.008; g = 1.90). FITC-dextran extravasation area was also larger in IsO2 PT: 13.42 ± 4.03 mm² compared with 6.60 ± 2.94 mm² in Awake PT (n = 6 and 5, respectively; Figure 10C). The between-group difference was 6.82 mm² (95% CI, 2.04–11.60; Welch’s t-test, p = 0.010; g = 1.74).

To account for differences in lesion size, the ratio of extravasation area to WFOI-derived lesion area at 1.5 h was also calculated for each animal and expressed as a percentage. The normalized value was 94.45 ± 33.97% in Awake PT (n = 5) and 167.24 ± 75.97% in IsO_2_ PT (n = 6). A large standardized between-group effect size was retained after normalization (difference, 72.78 percentage points; 95% CI, −8.48 to 154.05; Welch’s t-test, p = 0.072; g = 1.09).

Thus, photothrombosis under isoflurane anesthesia with oxygen as the carrier gas was associated with larger absolute areas of impaired capillary perfusion and dextran extravasation. The numerical difference in extravasation area between the two photothrombosis groups persisted even when expressed relative to the WFOI-derived lesion area.

### Photothrombosis under Isoflurane Anesthesia with Oxygen Is Associated with a More Prolonged Hemodynamic Response to Spreading Depolarization

During the 3–15 min period after injury, WFOI recordings revealed spreading depolarizations (SDs) accompanied by changes in Δ[HbT] propagating across the cortex (Figure 11A). At least one SD was detected in 1 of 5 Laser animals, 4 of 5 Awake PT animals, and all 7 IsO_2_ PT animals. The mean number of recorded SDs during the 12-min observation period was 0.20 ± 0.45 in Laser (n = 5), 2.00 ± 1.87 in Awake PT (n = 5), and 2.43 ± 0.79 in IsO2 PT (n = 7). Event counts differed among groups (Poisson regression, p = 0.0016). More SDs were recorded in both photothrombosis groups than in Laser (IsO_2_ PT: p_adj_3 = 0.046; Awake PT: p_adj_3 = 0.056), whereas the mean event counts were comparable between Awake PT and IsO_2_ PT (p_adj_3 = 0.626; Figure 11D). Holm’s correction was applied across the three pairwise comparisons.

**Figure 11.**
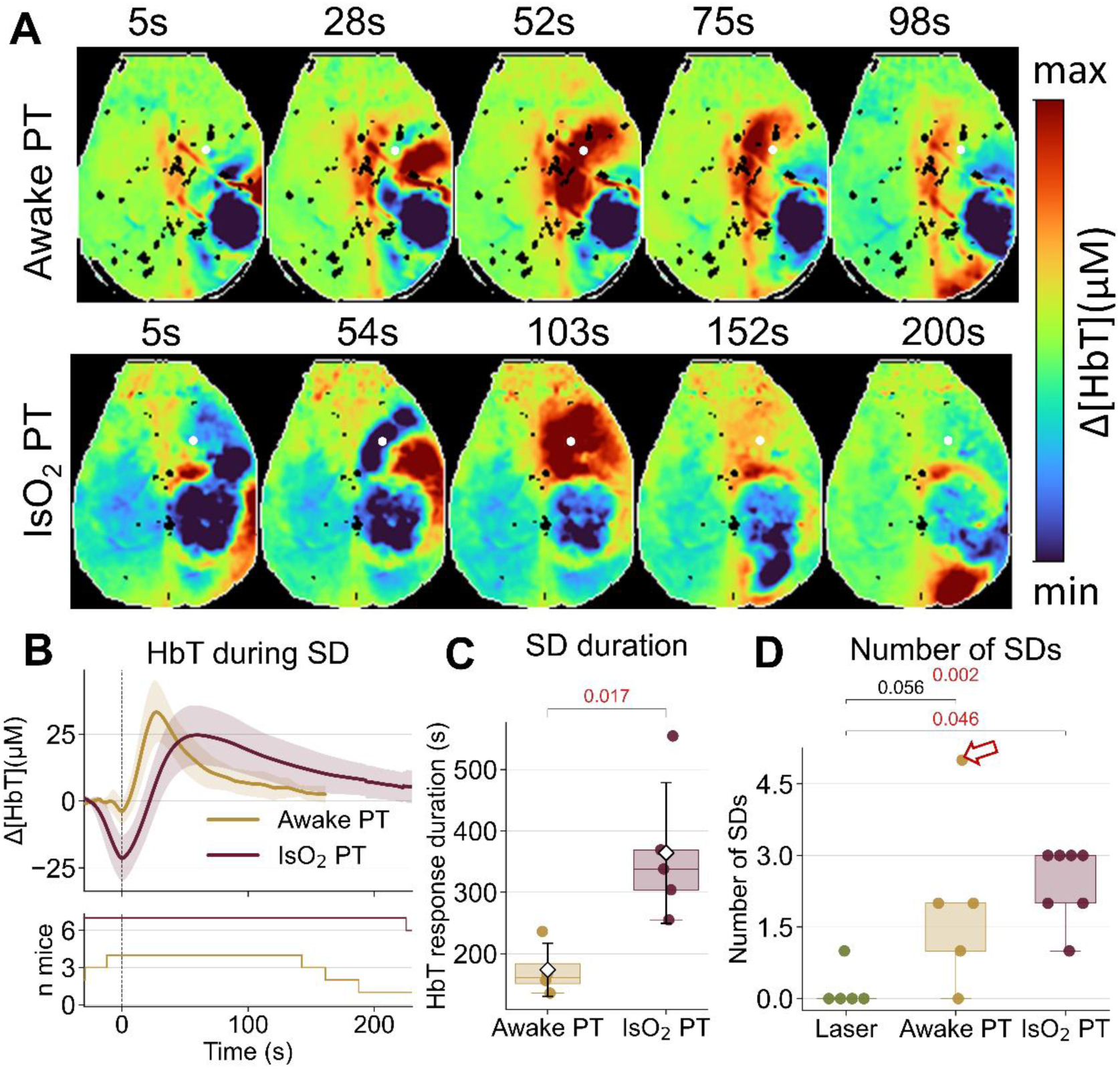
The hemodynamic response to spreading depolarization was longer after photothrombosis under isoflurane anesthesia. (A) Representative frames of hemodynamic changes accompanying a spreading depolarization (SD) in an awake mouse (top) and a mouse under isoflurane anesthesia (bottom). Pseudocolor indicates relative changes in total hemoglobin concentration (Δ[HbT]). Times are given in seconds from the onset of the HbT response. White circles indicate the regions from which HbT signals were extracted to assess the time course and duration of the hemodynamic response. (B) Group-averaged temporal Δ[HbT] profiles during SDs in the perifocal region, aligned to the HbT level at response onset and to the time of the HbT minimum. Curves represent means, and shaded areas indicate mean ± std. The number of animals contributing to each time point is shown below. (C) Duration of the SD-associated HbT response in Awake PT (n = 4) and IsO₂ PT (n = 5). The p-value was obtained using a two-sided Welch’s t-test. (D) Number of recorded SDs in Laser (n = 5), Awake PT (n = 5), and IsO₂ PT (n = 7). The arrow identifies the animal with a re-entrant spiral wave of spreading depolarization. Group differences were assessed using Poisson regression (likelihood-ratio test), with Holm-adjusted Wald tests for pairwise comparisons.

The photothrombosis groups also differed in the duration of the hemodynamic response accompanying SDs. Averaged Δ[HbT] profiles had a similar overall shape, but changes persisted longer in IsO_2_ PT (Figure 11B). When several SDs were recorded in one animal, profiles were averaged within the animal before group averaging. The duration of the complete HbT response was 173.8 ± 43.5 s in Awake PT (n = 4) and 364.1 ± 114.6 s in IsO_2_ PT (n = 5; Figure 11C). The hemodynamic response was, on average, 190.4 s longer in IsO2 PT (95% CI, 50.0–330.7 s; Welch’s t-test, p = 0.017; g = 1.86). Duration analyses included only animals for which both the onset and offset of a complete HbT response could be identified; incomplete events were excluded.

Thus, SDs were more frequently recorded after photothrombosis than after the milder laser-induced injury. Although the two photothrombosis groups showed similar mean event counts during the observation period, the HbT response to an individual SD was approximately twice as long in IsO_2_ PT as in Awake PT.

### Spreading Depolarization Counts and Dextran Extravasation Are Associated with Cortical Lesion Area

The number of recorded SDs correlated positively with WFOI-derived lesion area 10 min after irradiation (Spearman’s ρ = 0.639; n = 17; 95% bootstrap CI, 0.185–0.875; p = 0.0058). Animals with larger lesions had more SDs recorded during the 3–15 min interval. This association suggests that differences in SD counts may be related to the severity of the initial injury.

FITC-dextran extravasation area at 1.5 h after photothrombosis correlated positively with TTC-negative lesion area on day 3 (Spearman’s ρ = 0.867; n = 10; 95% bootstrap CI, 0.444–1.000; p = 0.0012). The number of recorded SDs also correlated positively with TTC-negative lesion area (ρ = 0.782; n = 15; 95% bootstrap CI, 0.340–0.956; p < 0.001).

Thus, the number of recorded SDs was associated both with WFOI-derived lesion area at 10 min and with TTC-negative lesion area on day 3; early dextran extravasation was also associated with the latter. These results indicate a close relationship between early vascular and functional disturbances and subsequent cortical injury severity.

## Discussion

### Isoflurane Anesthesia with Oxygen Enhances Slow Hemodynamic Oscillations at 0.05–0.10 Hz

Spontaneous cortical hemodynamic oscillations have diverse physiological origins and reflect not only changes in neuronal activity but also vasomotor and systemic vascular processes (Ma et al., 2016b; Mateo et al., 2017; Bumstead et al., 2017). Infra-slow activity (<0.08–0.10 Hz) exhibits pronounced spatiotemporal organization and can propagate across the cortex as large-scale waves (Matsui et al., 2016; Mitra et al., 2018). Hemodynamic oscillations in the 0.08–0.40 Hz range are more closely associated with changes in neuronal activity, although vasomotor and systemic vascular processes contribute alongside neurovascular coupling (Ma et al., 2016b; Mateo et al., 2017; Bumstead et al., 2017). At higher frequencies, correspondence between hemodynamic signals and neuronal activity decreases, whereas contributions from other physiological oscillations increase (Ma et al., 2016b; Wright et al., 2017; Turley et al., 2017).

In the present study, isoflurane anesthesia was associated with frequency-dependent reorganization of spontaneous hemodynamic activity. The most prominent change was an increase in relative spectral power at 0.05–0.10 Hz, predominantly in the somatosensory and primary motor cortices. This was consistent with the visually observed emergence of recurrent, widespread bilateral changes in the hemodynamic signal. Similar patterns have previously been described during simultaneous Ca²⁺ and hemodynamic imaging under anesthesia, in which propagating neuronal events were accompanied by spatially similar but delayed hemodynamic responses (Matsui et al., 2016).

Propagating cortical spatiotemporal patterns have also been reported for infra-slow activity, with characteristics depending on wakefulness or anesthesia (Mitra et al., 2018).

### Isoflurane Anesthesia Increases MOp Homotopic Connectivity and Alters RSP Functional Interactions

Functional connectivity (FC) describes the degree to which activity changes in different brain regions are coordinated. FC is widely studied using resting-state fMRI (Van Den Heuvel and Hulshoff Pol, 2010; White et al., 2011; Kura et al., 2018). Here, FC was determined from Pearson correlations between spontaneous hemodynamic signals in cortical regions (Song et al., 2011; White et al., 2011). Homotopic connectivity (HC) characterizes interhemispheric coordination between symmetrical cortical locations, supported in part by transcallosal connections (Vanni and Murphy, 2014; Aykan et al., 2021), whereas global brain connectivity (GBC) reflects the degree of concordance between each location and the remaining analyzed cortex (Cole et al., 2010). Interregional correlation matrices provide an additional measure of concordance in hemodynamic fluctuations between specific cortical regions. Importantly, FC is not a direct measure of anatomical connectivity.

In our study, changes in FC under isoflurane anesthesia were primarily regional. HC increased in MOp, whereas correlations between RSP hemodynamic fluctuations and those in other analyzed regions decreased. The increased HC in MOp is consistent with enhanced bilateral synchronization of slow hemodynamic activity in the sensorimotor cortex. In contrast, the reduced correlations involving RSP suggest a relative functional segregation of this region from the motor and somatosensory cortices.

The effects of anesthesia on cortical functional organization have also been demonstrated in previous studies, although their direction depends on the experimental conditions. In a mouse fMRI study, increasing isoflurane concentrations reduced homotopic connectivity and disrupted the modular organization of functional networks (Bukhari et al., 2018), in contrast to the regional increase in homotopic connectivity observed in our study. Wide-field optical imaging has also revealed differences in spatial patterns and global connectivity among several anesthetic regimens, including isoflurane (Xie et al., 2020). The particular involvement of RSP is supported by Ca²⁺ imaging data: deeper isoflurane anesthesia increased the representation of retrosplenial activity components and correlations among them (Scaglione et al., 2024). These findings demonstrate that anesthesia may differentially affect the functional organization of individual brain regions and networks. Under our experimental conditions, the effects of anesthesia on functional connectivity were region-specific, suggesting differential sensitivity of cortical regions to anesthetic conditions.

### Induction of Isoflurane Anesthesia in Oxygen Increases Cortical Blood Oxygenation

The additional experiment showed that induction of isoflurane anesthesia using 100% O₂ was accompanied by increased oxygenated (HbO) and total (HbT) hemoglobin concentrations and decreased deoxygenated hemoglobin (HHb), indicating increased cortical blood oxygenation and consistent with an increased blood volume within the analyzed tissue. The increase in HbT may be related to the vasodilatory action of isoflurane. In mice, the transition from wakefulness to isoflurane anesthesia has been associated with cerebral vasodilation and increased cerebral blood flow (Sullender et al., 2022; Pan et al., 2023), as well as increased HbT, including when air was used as the carrier gas (Qin et al., 2025). Conversely, hyperoxia may cause cerebral vasoconstriction and reduced cerebral blood flow (Pilkinton et al., 2012), although such a decrease was not observed in spontaneously breathing rats under isoflurane anesthesia, potentially because of a concurrent increase in arterial partial pressure of CO₂ (PaCO₂) (Sicard and Duong, 2005). Furthermore, 100% O₂ increases cortical blood oxygenation but does not necessarily increase HbT (Sharp et al., 2015). Elevated cerebral blood oxygenation may also reflect an isoflurane-induced reduction in the cerebral metabolic rate of oxygen (CMRO₂), consistent with decreased tissue oxygen consumption (Cao et al., 2017).

Thus, the observed changes are consistent with isoflurane-induced vasodilation and increased cortical blood oxygenation, potentially accompanied by reduced cerebral oxygen consumption. These findings show that the conditions of anesthesia induction substantially alter the baseline cortical hemodynamic state and may consequently modify the conditions under which photothrombotic injury develops.

### Laser Exposure without Effective Intravenous Photosensitizer Administration Also Produced Cortical Injury

In the Laser group, in which effective intravenous delivery of Rose Bengal could not be achieved, focal cortical injury nevertheless occurred, including the development of a TTC-negative lesion on day 3. This suggests that injury may have arisen not only from the classical photochemical reaction but also from direct effects of laser irradiation.

Direct laser exposure may have contributed to the central portion of the lesion, whereas effective photosensitizer administration may have produced additional injury in the surrounding tissue through a photochemical mechanism (Watson et al., 1985). However, such a spatial separation of injury mechanisms was not directly investigated in the present study.

Classical photothrombosis protocols use lasers with wavelengths of approximately 532–568 nm and powers ranging from a few to several tens of milliwatts (Sadanaga et al., 1998; García-Peña et al., 2023). Nevertheless, protocols employing higher laser powers, up to 400 mW, have also been reported (Wang et al., 2026). The nominal laser power in our experiment was approximately 1000 mW. Tissue heating may also occur at much lower illumination intensities (Chary et al., 2026). Direct laser-mediated vascular injury has additionally been reported using femtosecond pulses (Luo et al., 2017), although this irradiation regime differs substantially from ours.

It should also be considered that the actual laser output may differ from the manufacturer’s nominal specification and vary during irradiation, thereby affecting lesion-size reproducibility (Alaverdashvili et al., 2015). Because neither the power delivered to the cortical surface nor tissue temperature was measured directly, the contribution of photothermal effects cannot be estimated. In addition, some Rose Bengal may have entered the circulation after subcutaneous delivery. Thus, Laser was the group with the mildest injury, but it cannot be considered a strict laser-only control in the complete absence of photosensitizer.

### Differences in Cortical Injury Severity Were Established during Injury Induction

Differences in cortical lesion area between experimental conditions were detectable within 10 min after irradiation. The largest lesions occurred in IsO_2_ PT, where lesion area was significantly greater than in Awake PT and Laser. These findings indicate that differences in injury severity were associated with the conditions of induction and were already evident in the hyperacute period, rather than arising solely through subsequent pathological progression. By day 3, TTC staining demonstrated a graded severity pattern, Laser < Awake PT < IsO_2_ PT, with statistically significant differences among all three groups.

Longitudinal WFOI assessment revealed different lesion-area trajectories. Lesion area increased in all groups by day 1, but between days 1 and 3 it decreased in Laser, remained nearly unchanged in Awake PT, and continued to increase in IsO^2^ PT. Similar dynamics have been described after experimental photothrombosis, where changes in lesion size during the first 1–3 days may reflect not only the spread of irreversible injury but also tissue edema (Li et al., 2014; Liu et al., 2017). Thus, differences in final injury severity may have been determined both by the magnitude of the initial insult and by its subsequent evolution. WFOI-derived lesion area on day 3 correlated strongly with TTC-negative tissue area, consistent with previous demonstrations of optical mapping of ischemic injury (Sunil et al., 2021).

Differences between induction conditions were also evident in early functional outcomes. HC decreased in all three groups as early as 1.5 h after injury, including in Laser, but the greatest decrease occurred in IsO_2_ PT. Alterations involved not only SS and the adjacent MOp, corresponding to the main lesion region, but also the less affected MOs and RSP, where early between-group differences were detected. Pronounced regional heterogeneity of homotopic connectivity disturbances following focal ischemia has also been described previously (Bauer et al., 2014; Hakon et al., 2018; Cramer et al., 2019). Notably, the early HC reduction in MOs was closely associated with TTC-negative lesion area on day 3. By contrast, changes in GBC within the intact hemisphere were less pronounced, with a transient numerical increase on day 1 most evident in Awake PT.

Early vascular disturbances were likewise most pronounced in IsO_2_ PT. At 1.5 h after photothrombosis, this group had larger areas of impaired capillary perfusion and FITC-dextran extravasation than Awake PT. Early BBB permeability disruption after photothrombotic ischemia is consistent with two-photon imaging studies (Underly et al., 2017; Underly and Shih, 2017; Protzmann et al., 2024). Extravasation area at 1.5 h was strongly correlated with TTC-negative lesion area on day 3. However, even after normalization to the lesion area measured by WFOI at 1.5 h after photothrombosis, the extravasation-to-lesion area ratio remained higher in IsO_2_ PT than in Awake PT, with a large standardized effect size (g = 1.09; p = 0.072). This suggests that the extent of BBB disruption may depend not only on lesion size but also on the conditions of photothrombosis induction.

Independent motor and biochemical outcomes also supported differences in injury severity, although their sensitivity varied. The Grid test detected persistent motor deficits even after relatively mild injury in Laser, whereas the Cylinder test and NSE concentrations more clearly differentiated injury severity across the experimental conditions. The increase in NSE by day 3 is consistent with its use as a biomarker of neuronal injury in cerebral ischemia (Gelderblom et al., 2013; Isgrò et al., 2015; Mochetti et al., 2024). Furthermore, increased Grid-test errors and NSE levels were closely associated with final TTC-negative lesion area.

Thus, differences among experimental conditions emerged within minutes of irradiation and were subsequently reflected in the severity of vascular and functional disturbances, final lesion size, and motor and biochemical outcomes. This indicates that the conditions under which injury was induced played a major role in determining the severity of cortical injury in our study.

### Isoflurane Anesthesia with Oxygen Prolongs the Hemodynamic Response to Spreading Depolarization

Spreading depolarizations (SDs) were another early manifestation of injury. They were recorded more frequently after photothrombosis than after the relatively mild laser-induced injury, and higher recorded SD counts were associated with both WFOI-derived lesion area at 10 min and final TTC-negative lesion area. While mean event counts were similar between the two photothrombosis groups, the HbT response to an individual SD was approximately twice as long in IsO2 PT as in Awake PT.

In several experimental models, isoflurane has been associated with lower SD frequency compared with other anesthetic regimens or with suppression of SD initiation, depending on the experimental conditions (Klass et al., 2018; Kudo et al., 2008; Takagaki et al., 2014). Conversely, direct comparisons between awake and anesthetized mice showed longer-lasting SDs under anesthesia despite comparable induction thresholds and propagation velocities (Houben et al., 2017). Nevertheless, studies directly comparing SD characteristics in awake and anesthetized animals remain limited (Sonn and Mayevsky, 2006; Houben et al., 2017; Boyce et al., 2025).

Unlike the other functional outcomes, the SD-associated hemodynamic response in IsO_2_ PT was recorded directly under anesthesia. Importantly, isoflurane itself modifies baseline cerebral hemodynamics and neurovascular coupling, including cerebral blood flow and the temporal properties of hemodynamic responses (Masamoto et al., 2009; Pisauro et al., 2013; Cao et al., 2017). In addition, the use of oxygen as the carrier gas and spontaneous breathing under anesthesia may have altered vascular tone independently of the direct effects of isoflurane. Hyperoxia, for example, may cause cerebral vasoconstriction, whereas CO₂ accumulation due to reduced ventilation may produce vasodilation (Floyd et al., 2003; Sicard and Duong, 2005). It should be noted, however, that arterial CO₂ partial pressure was not monitored in our experiment, and the effects of isoflurane and the carrier gas were not independently controlled.

The measured duration of the HbT response characterizes the hemodynamic manifestation of an SD rather than the duration of the underlying electrophysiological depolarization. The prolonged HbT response in IsO2 PT may reflect altered vascular reactivity associated with isoflurane anesthesia and hyperoxia, as well as greater injury severity. However, in accordance with previously published results, we assume a prevailing effect of isoflurane (Houben et al., 2017).

The recorded events most likely represent peri-infarct depolarizations occurring during the early post-irradiation period. As recordings covered a limited observation window, the earliest and later SDs may not have been captured, and prolonged hemodynamic responses may have hindered the separation of successive events. Nevertheless, the pronounced prolongation of the HbT response in IsO_2_ PT indicates that anesthetic conditions are associated with altered hemodynamic manifestations of SDs during the acute phase of cortical injury.

### Isoflurane Anesthesia in an Oxygen-Rich Gas Mixture Was Associated with More Severe Injury

In the present study, photothrombosis induced under isoflurane anesthesia with oxygen as the carrier gas resulted in the most severe structural, vascular, and functional consequences. This contrasts with several experimental studies reporting neuroprotective effects of isoflurane in cerebral ischemia, including reduced infarct volume after middle cerebral artery occlusion or photothrombotic injury (Kapinya et al., 2002; Seto et al., 2014; Lu et al., 2017; Boyce et al., 2025). Proposed mechanisms of isoflurane-mediated neuroprotection include reduced cerebral metabolic demands, attenuation of excitotoxicity and inflammation, and activation of cellular survival pathways (Wang et al., 2016; Archer et al., 2017).

An important feature of our experimental protocol was the use of oxygen as the carrier gas. Notably, in the photothrombosis model, the neuroprotective effect of isoflurane was demonstrated using air as the carrier gas (Seto et al., 2014). However, the composition of the inhaled gas mixture during photothrombosis induction is not always explicitly reported (Lu et al., 2017; Boyce et al., 2025). The mechanism of Rose Bengal-mediated photothrombosis is directly dependent on molecular oxygen: photoactivation of the dye produces singlet oxygen and other reactive oxygen species that damage the vascular endothelium and trigger thrombosis (Saniabadi et al., 1995; Uzdensky, 2018). Breathing pure oxygen has previously been shown to accelerate photochemical occlusion of retinal vessels after Rose Bengal administration (Wilson and Hatchell, 1991). Thus, increased oxygen availability could potentially have enhanced photochemical vascular damage.

In our additional experiment, induction of isoflurane anesthesia in oxygen was associated with increased cortical blood oxygenation. The increased cortical blood oxygenation may have facilitated oxygen-dependent photochemical reactions within cerebral vessels. Consistent with this hypothesis, more extensive injury was detected in IsO_2_ PT within 10 min of irradiation, indicating that the differences between induction conditions arose early.

Isoflurane and hyperoxia may also affect cerebral hemodynamics and vascular tone. Isoflurane-induced cerebral vasodilation may increase cerebral blood volume (Sullender et al., 2022; Qin et al., 2025). Hyperoxia has been associated with cerebral vasoconstriction and decreased cerebral blood flow (Floyd et al., 2003; Lu et al., 2009). Altered gas exchange, including absorption atelectasis during breathing of pure oxygen, may represent an additional factor (Wilding et al., 2017).

Thus, more severe injury in IsO_2_ PT may have resulted not from isoflurane alone but from its combination with a high inspired oxygen concentration, which could enhance the photochemical reaction and alter the vascular conditions of lesion formation. These findings emphasize that both the choice of anesthetic and the composition of the inhaled gas mixture should be considered when standardizing photothrombotic models.

### Study Limitations

This study has several limitations. Small experimental group sizes, further reduced in some analyses by animal mortality and technical difficulties, limited statistical power, particularly for between-group and regional comparisons. In addition, the Laser group was formed post hoc rather than as a prospectively planned randomized control group.

The injury-induction protocol did not allow the separate effects of isoflurane and oxygen to be determined. The absence of direct measurements of tissue pO₂, CO₂, and cortical temperature further limits interpretation of the mechanisms potentially responsible for exacerbated injury. Actual laser power at the tissue surface was also not measured.

Spectral analysis was constrained by the duration of available quiet-wakefulness periods: 60-s recordings ensured comparability but reduced the reliability of estimates at the lowest frequencies. Formal blinding was not used during data processing, and some measures were delineated manually, potentially introducing operator-dependent variability. Microcirculation and BBB permeability were assessed only in the two photothrombosis groups.

### Conclusions

Using WFOI, we demonstrated that anesthetic conditions influence both hemodynamic activity in the intact cortex and the severity of injury induced by photothrombosis. Isoflurane anesthesia under hyperoxic conditions was associated with increased relative power of slow hemodynamic oscillations and altered regional functional connectivity. Differences in lesion severity between experimental conditions were detectable within 10 min after irradiation, and TTC staining confirmed a graded increase in injury severity by day 3. The most severe structural, vascular, and functional disturbances occurred when photothrombosis was induced under isoflurane anesthesia with oxygen as the carrier gas.

The observed increase in cortical blood oxygenation during anesthesia induction suggests that oxygen-dependent photochemical processes may contribute to injury exacerbation. Together, these findings demonstrate the utility of WFOI for assessing cortical injury from its earliest stages and highlight the importance of anesthetic conditions, including the composition of the inhaled gas mixture, in the standardization of experimental stroke models and the interpretation of cortical hemodynamic activity.

## Supplementary Materials

Tables S1–S5: Detailed descriptive statistics and statistical comparisons.

Video S1: Comparison of spontaneous cortical hemodynamic activity (Δ[HbT]) during wakefulness and isoflurane anesthesia.

## Author Contributions

Lizunova N.V. Investigation, Methodology, Conceptualization, Data curation, Writing – Original Draft, Formal Analysis, Software, Visualization

Kislukhina E.N. Investigation, Methodology, Conceptualization, Data curation, Writing – Original Draft, Formal Analysis

Dubas V.I. Investigation, Formal Analysis Alekseev V.I. Writing – Review & Editing

Surin A.M. Writing – Review & Editing, Supervision

Bakaeva Z.V. Conceptualization, Writing – Review & Editing, Supervision

## Funding

This research was funded by the Russian Science Foundation (RSF), grant № 25-25-00727, https://rscf.ru/en/project/25-25-00727/.

## Institutional Review Board Statement

The animal study protocol was approved by the Ethics Committee of the National Medical Research Center for Children’s Health (Protocol No. 1, 30 January 2018). All animal procedures were conducted in accordance with Directive 2010/63/EU.

## Informed Consent Statement

Not applicable.

## Data Availability Statement

The data presented in this study are available from the corresponding author upon reasonable request. The code used for WFOI data processing, statistical analysis, and figure generation is available at https://github.com/natalia1896/WIFIOPIA_PT2021_git.

## Acknowledgments

The authors would like to thank Vladislav Kuzmin’s research group, particularly Alexandra Ivanova, for providing access to the isoflurane anesthesia system.

## Conflicts of Interest

The authors declare no conflicts of interest. The funder had no role in the design of the study; in the collection, analyses, or interpretation of data; in the writing of the manuscript; or in the decision to publish the results.

## Abbreviations

The following abbreviations are used in this manuscript:

ANOVA: Analysis of variance
BBB: Blood–brain barrier
CI: Confidence interval
CMRO₂: Cerebral metabolic rate of oxygen
ELISA: Enzyme-linked immunosorbent assay
FC: Functional connectivity
FITC: Fluorescein isothiocyanate
fMRI: Functional magnetic resonance imaging
fNIRS: Functional near-infrared spectroscopy
GBC: Global brain connectivity
HbO: Oxygenated hemoglobin
HbT: Total hemoglobin
HC: Homotopic connectivity
HHb: Deoxygenated hemoglobin
IQR: Interquartile range
MOp: Primary motor cortex
Mos: Secondary motor cortex
NSE: Neuron-specific enolase
OISI: Optical intrinsic signal imaging
PaCO₂: Arterial partial pressure of carbon dioxide
pp: Percentage points
PSD: Power spectral density
PT: Photothrombosis
ROI: Region of interest
RSP: Retrosplenial cortex
SD: Spreading depolarization
sem: Standard error of the mean
SS: Somatosensory cortex
std: Standard deviation
TTC: 2,3,5-Triphenyltetrazolium chloride
WFOI: Wide-field optical imaging

## Notes

### Competing Interest Statement

The authors have declared no competing interest.

