## Supplementary Tables S1S5 for "Isoflurane Anesthesia under Hyperoxic Conditions Shapes Cortical Hemodynamics and Photothrombotic Injury Severity in Mice: A Wide-Field Optical Imaging Study"

**Table S1. Spontaneous Hemodynamic Activity and Functional Connectivity during Wakefulness and Isoflurane Anesthesia**

| **Measure** | **Scope / ROI** | **Awake, mean ± std (n)** | **Iso, mean ± std (n)** | **Δ Iso − Awake** | **95% CI** | **t** | **df** | **p_raw** | **p_adj** | **Hedges g** |
| --- | --- | --- | --- | --- | --- | --- | --- | --- | --- | --- |
| Relative spectral power  0.05–0.10 Hz | Whole brain | 25,89 ± 9,81 (9) | 38,27 ± 9,28 (7) | +12,38 | 2,04…22,71 | 2,58 | 13,38 | **0,022** | — | +1,22 |
|  | MOp | 22,46 ± 12,18 (9) | 39,07 ± 10,07 (7) | +16,61 | 4,66…28,55 | 2,98 | 13,91 | **0,010** | **0,030** | +1,39 |
|  | MOs | 27,72 ± 9,76 (9) | 39,79 ± 11,30 (7) | +12,08 | 0,37…23,78 | 2,25 | 11,96 | **0,044** | **0,088** | +1,09 |
|  | SS | 23,33 ± 10,08 (9) | 37,16 ± 6,28 (7) | +13,83 | 4,97…22,68 | 3,36 | 13,49 | **0,005** | **0,019** | +1,51 |
|  | RSP | 25,73 ± 11,55 (8) | 29,66 ± 10,03 (6) | +3,93 | −8,71…16,57 | 0,68 | 11,66 | 0,510 | 0,510 | +0,34 |
| Relative spectral power 0.11–0.20 Hz | Whole brain | 31,33 ± 7,43 (9) | 30,19 ± 9,13 (7) | −1,14 | −10,44…8,16 | −0,27 | 11,49 | 0,794 | — | −0,13 |
|  | MOp | 32,59 ± 7,66 (9) | 30,93 ± 9,08 (7) | −1,66 | −11,00…7,68 | −0,39 | 11,78 | 0,705 | 1,000 | −0,19 |
|  | MOs | 30,51 ± 9,46 (9) | 27,99 ± 7,79 (7) | −2,52 | −11,78…6,74 | −0,58 | 13,92 | 0,569 | 1,000 | −0,27 |
|  | SS | 31,95 ± 6,71 (9) | 31,67 ± 10,36 (7) | −0,28 | −10,36…9,81 | −0,06 | 9,77 | 0,953 | 1,000 | −0,03 |
|  | RSP | 32,47 ± 10,24 (8) | 31,70 ± 10,54 (6) | −0,77 | −13,18…11,65 | −0,14 | 10,74 | 0,894 | 1,000 | −0,07 |
| Relative spectral power  0.21–0.40 Hz | Whole brain | 26,03 ± 3,75 (9) | 18,53 ± 8,49 (7) | −7,50 | −15,48…0,47 | −2,18 | 7,83 | **0,062** | — | −1,14 |
|  | MOp | 27,14 ± 6,40 (9) | 17,57 ± 9,60 (7) | −9,57 | −18,94…−0,19 | −2,27 | 9,97 | **0,046** | 0,185 | −1,14 |
|  | MOs | 25,88 ± 3,23 (9) | 19,08 ± 7,84 (7) | −6,81 | −14,15…0,54 | −2,16 | 7,59 | **0,065** | 0,185 | −1,13 |
|  | SS | 27,06 ± 7,92 (9) | 18,01 ± 9,19 (7) | −9,04 | −18,56…0,47 | −2,07 | 11,95 | **0,061** | 0,185 | −1,01 |
|  | RSP | 24,01 ± 6,80 (8) | 23,77 ± 10,45 (6) | −0,24 | −11,52…11,03 | −0,05 | 8,10 | 0,962 | 0,962 | −0,03 |
| Homotopic functional connectivity (HC) | Pixelwise whole cortex | 0,771 ± 0,089 (9) / 1,068 ± 0,270 (9) | 0,826 ± 0,125 (7) / 1,268 ± 0,366 (7) | +0,200 | −0,165…0,564 | 1,21 | 10,72 | 0,253 | — | +0,60 |
|  | MOp | 0,786 ± 0,093 (9) / 1,105 ± 0,252 (9) | 0,923 ± 0,055 (7) / 1,706 ± 0,337 (7) | +0,601 | 0,265…0,937 | 3,94 | 10,84 | **0,002** | **0,010** | +1,95 |
|  | MOs | 0,729 ± 0,124 (9) / 0,989 ± 0,328 (9) | 0,742 ± 0,177 (7) / 1,054 ± 0,410 (7) | +0,065 | −0,351…0,481 | 0,34 | 11,35 | 0,739 | 1,000 | +0,17 |
|  | SS | 0,837 ± 0,085 (8) / 1,300 ± 0,378 (8) | 0,866 ± 0,112 (6) / 1,459 ± 0,446 (6) | +0,159 | −0,345…0,664 | 0,71 | 9,80 | 0,497 | 1,000 | +0,37 |
|  | RSP | 0,837 ± 0,087 (8) / 1,287 ± 0,336 (8) | 0,832 ± 0,117 (6) / 1,314 ± 0,429 (6) | +0,027 | −0,450…0,504 | 0,13 | 9,26 | 0,901 | 1,000 | +0,07 |
| Global functional connectivity (GBC) | Pixelwise bilateral cortex | 0,493 ± 0,046 (9) / 0,541 ± 0,058 (9) | 0,415 ± 0,203 (7) / 0,469 ± 0,276 (7) | −0,072 | −0,327…0,184 | −0,68 | 6,42 | 0,523 | — | −0,36 |
|  | MOp | 0,544 ± 0,031 (9) / 0,611 ± 0,043 (9) | 0,524 ± 0,255 (7) / 0,641 ± 0,369 (7) | +0,030 | −0,312…0,371 | 0,21 | 6,13 | 0,838 | 1,000 | +0,12 |
|  | MOs | 0,455 ± 0,066 (9) / 0,493 ± 0,080 (9) | 0,395 ± 0,196 (7) / 0,441 ± 0,257 (7) | −0,052 | −0,291…0,186 | −0,52 | 6,92 | 0,619 | 1,000 | −0,28 |
|  | SS | 0,552 ± 0,084 (9) / 0,629 ± 0,123 (9) | 0,526 ± 0,223 (7) / 0,635 ± 0,339 (7) | +0,006 | −0,310…0,322 | 0,04 | 7,23 | 0,966 | 1,000 | +0,02 |
|  | RSP | 0,524 ± 0,079 (8) / 0,588 ± 0,113 (8) | 0,240 ± 0,309 (6) / 0,273 ± 0,361 (6) | −0,315 | −0,692…0,063 | −2,06 | 5,74 | **0,087** | 0,347 | −1,19 |

**Note.** For spectral power, values are percentages and Δ is expressed in percentage points. The three bands shown are 0.05–0.10, 0.11–0.20, and 0.21–0.40 Hz. For HC and GBC, group values are Pearson r / Fisher z; differences, 95% CIs, t and g refer to Fisher z. Regional p_adj values use Holm correction across MOp/MOs/SS/RSP; scalar whole-brain/composite/pixelwise rows are not regionally adjusted (—). MOp, primary motor cortex; MOs, secondary motor cortex; SS, somatosensory cortex; RSP, retrosplenial cortex. Values with p < 0.10 are bold burgundy; |g| ≥ 0.80 is burgundy.

**Table S2. Regional and Pixelwise Changes in Homotopic Functional Connectivity after Focal Cortical Injury**

| **Scope / ROI** | **Contrast** | **Laser, Δz mean ± SD (n) 95% CI** | **p_vs0 raw** | **p_vs0 adj** | **Awake PT, Δz mean ± SD (n) 95% CI** | **p_vs0 raw** | **p_vs0 adj** | **Iso PT, Δz mean ± SD (n) 95% CI** | **p_vs0 raw** | **p_vs0 adj** | **Welch F** | **df1** | **df2** | **p_group raw** | **p_group adj** |
| --- | --- | --- | --- | --- | --- | --- | --- | --- | --- | --- | --- | --- | --- | --- | --- |
| Pixelwise whole cortex | 1.5 h − Baseline | −0.317 ± 0.135 (5) 95% CI −0.485…−0.150 | **0.006** | **0.013** | −0.633 ± 0.311 (5) 95% CI −1.019…−0.247 | **0.010** | **0.013** | −0.881 ± 0.179 (7) 95% CI −1.046…−0.716 | **<0.001** | **<0.001** | 17.95 | 2 | 8.18 | **0.001** | — |
|  | Day 1 − Baseline | −0.430 ± 0.348 (5) 95% CI −0.862…0.002 | **0.051** | **0.059** | −0.584 ± 0.394 (5) 95% CI −1.073…−0.095 | **0.029** | **0.059** | −0.857 ± 0.101 (5) 95% CI −0.983…−0.731 | **<0.001** | **<0.001** | 3.91 | 2 | 6.05 | **0.081** | — |
|  | Day 3 − Day 1 | −0.041 ± 0.077 (4) 95% CI −0.163…0.081 | 0.365 | 1.000 | 0.075 ± 0.324 (5) 95% CI −0.327…0.477 | 0.632 | 1.000 | 0.093 ± 0.229 (5) 95% CI −0.191…0.377 | 0.414 | 1.000 | 0.88 | 2 | 6.24 | 0.461 | — |
|  | Day 3 − Baseline | −0.465 ± 0.308 (5) 95% CI −0.847…−0.083 | **0.028** | **0.028** | −0.509 ± 0.142 (5) 95% CI −0.686…−0.333 | **0.001** | **0.004** | −0.764 ± 0.243 (5) 95% CI −1.065…−0.462 | **0.002** | **0.004** | 2.12 | 2 | 7.20 | 0.189 | — |
| MOp | 1.5 h − Baseline | −0.696 ± 0.429 (5) 95% CI −1.229…−0.164 | **0.022** | **0.022** | −1.063 ± 0.417 (5) 95% CI −1.581…−0.545 | **0.005** | **0.009** | −1.220 ± 0.246 (7) 95% CI −1.448…−0.993 | **<0.001** | **<0.001** | 2.80 | 2 | 7.25 | 0.125 | 0.251 |
|  | Day 1 − Baseline | −0.792 ± 0.463 (5) 95% CI −1.367…−0.216 | **0.019** | **0.019** | −1.308 ± 0.616 (5) 95% CI −2.073…−0.543 | **0.009** | **0.018** | −1.314 ± 0.233 (5) 95% CI −1.603…−1.025 | **<0.001** | **<0.001** | 2.37 | 2 | 6.85 | 0.165 | 0.496 |
|  | Day 3 − Day 1 | 0.129 ± 0.084 (4) 95% CI −0.005…0.262 | **0.054** | 0.163 | 0.227 ± 0.463 (5) 95% CI −0.348…0.802 | 0.335 | 0.670 | −0.056 ± 0.321 (5) 95% CI −0.455…0.342 | 0.715 | 0.715 | 0.82 | 2 | 5.93 | 0.486 | 1.000 |
|  | Day 3 − Baseline | −0.623 ± 0.413 (5) 95% CI −1.136…−0.110 | **0.028** | **0.028** | −1.082 ± 0.306 (5) 95% CI −1.461…−0.702 | **0.001** | **0.004** | −1.370 ± 0.388 (5) 95% CI −1.852…−0.889 | **0.001** | **0.004** | 4.06 | 2 | 7.85 | **0.062** | 0.246 |
| MOs | 1.5 h − Baseline | −0.146 ± 0.124 (5) 95% CI −0.300…0.008 | **0.058** | **0.093** | −0.421 ± 0.331 (5) 95% CI −0.832…−0.010 | **0.047** | **0.093** | −0.680 ± 0.167 (7) 95% CI −0.835…−0.525 | **<0.001** | **<0.001** | 18.62 | 2 | 8.06 | **<0.001** | **0.004** |
|  | Day 1 − Baseline | −0.175 ± 0.384 (5) 95% CI −0.651…0.302 | 0.366 | 0.558 | −0.198 ± 0.353 (5) 95% CI −0.636…0.241 | 0.279 | 0.558 | −0.554 ± 0.127 (5) 95% CI −0.711…−0.397 | **<0.001** | **0.002** | 3.65 | 2 | 6.38 | **0.088** | 0.350 |
|  | Day 3 − Day 1 | −0.201 ± 0.294 (4) 95% CI −0.668…0.266 | 0.265 | 0.604 | −0.036 ± 0.364 (5) 95% CI −0.488…0.417 | 0.837 | 0.837 | 0.184 ± 0.269 (5) 95% CI −0.150…0.519 | 0.201 | 0.604 | 1.93 | 2 | 7.03 | 0.215 | 0.858 |
|  | Day 3 − Baseline | −0.351 ± 0.256 (5) 95% CI −0.670…−0.033 | **0.037** | **0.075** | −0.233 ± 0.226 (5) 95% CI −0.513…0.047 | **0.082** | **0.082** | −0.370 ± 0.180 (5) 95% CI −0.593…−0.147 | **0.010** | **0.030** | 0.55 | 2 | 7.82 | 0.598 | 1.000 |
| SS | 1.5 h − Baseline | −0.814 ± 0.641 (4) 95% CI −1.834…0.205 | **0.085** | **0.085** | −1.130 ± 0.228 (4) 95% CI −1.492…−0.768 | **0.002** | **0.004** | −1.439 ± 0.259 (5) 95% CI −1.760…−1.118 | **<0.001** | **<0.001** | 2.53 | 2 | 5.83 | 0.162 | 0.251 |
|  | Day 1 − Baseline | −1.207 ± 0.872 (4) 95% CI −2.593…0.180 | **0.070** | **0.070** | −1.542 ± 0.533 (5) 95% CI −2.204…−0.880 | **0.003** | **0.006** | −1.416 ± 0.157 (5) 95% CI −1.611…−1.221 | **<0.001** | **<0.001** | 0.22 | 2 | 5.00 | 0.808 | 0.808 |
|  | Day 3 − Day 1 | 0.322 ± 0.446 (3) 95% CI −0.786…1.431 | 0.337 | 0.981 | 0.258 ± 0.517 (5) 95% CI −0.384…0.901 | 0.327 | 0.981 | 0.026 ± 0.547 (4) 95% CI −0.843…0.896 | 0.929 | 0.981 | 0.31 | 2 | 5.40 | 0.749 | 1.000 |
|  | Day 3 − Baseline | −1.026 ± 0.621 (4) 95% CI −2.014…−0.037 | **0.046** | **0.046** | −1.284 ± 0.330 (5) 95% CI −1.693…−0.874 | **<0.001** | **0.003** | −1.353 ± 0.533 (4) 95% CI −2.201…−0.505 | **0.015** | **0.030** | 0.32 | 2 | 5.49 | 0.739 | 1.000 |
| RSP | 1.5 h − Baseline | −0.027 ± 0.244 (3) 95% CI −0.633…0.578 | 0.864 | 0.864 | −0.403 ± 0.188 (4) 95% CI −0.702…−0.103 | **0.024** | **0.047** | −0.860 ± 0.212 (7) 95% CI −1.056…−0.664 | **<0.001** | **<0.001** | 13.71 | 2 | 4.96 | **0.010** | **0.029** |
|  | Day 1 − Baseline | −0.402 ± 0.495 (3) 95% CI −1.631…0.828 | 0.295 | 0.295 | −0.521 ± 0.462 (5) 95% CI −1.095…0.053 | **0.065** | 0.131 | −0.822 ± 0.253 (5) 95% CI −1.136…−0.508 | **0.002** | **0.006** | 1.30 | 2 | 4.60 | 0.357 | 0.715 |
|  | Day 3 − Day 1 | −0.106 ± 0.099 (2) 95% CI −0.999…0.786 | 0.371 | 1.000 | 0.078 ± 0.409 (5) 95% CI −0.430…0.586 | 0.692 | 1.000 | 0.018 ± 0.404 (4) 95% CI −0.625…0.661 | 0.934 | 1.000 | 0.49 | 2 | 5.20 | 0.637 | 1.000 |
|  | Day 3 − Baseline | −0.489 ± 0.468 (3) 95% CI −1.651…0.673 | 0.212 | 0.212 | −0.443 ± 0.148 (5) 95% CI −0.627…−0.259 | **0.003** | **0.008** | −0.791 ± 0.393 (4) 95% CI −1.417…−0.166 | **0.028** | **0.055** | 1.19 | 2 | 3.67 | 0.399 | 1.000 |

**Note.** All changes are Fisher Δz. Contrasts: 1.5 h − baseline; day 1 − baseline; day 3 − day 1; day 3 − baseline. p_vs0 adj, Holm correction across the three experimental groups for one-sample comparisons with zero. For regional rows, p_group adj is Holm-corrected across the four ROIs; the pixelwise scalar has no regional multiplicity correction (—). Values with p < 0.10 are bold burgundy.

**Table S3. Regional and Pixelwise Changes in Global Functional Connectivity of the Intact Hemisphere after Focal Cortical Injury**

| **Scope / ROI** | **Contrast** | **Laser, Δz mean ± SD (n) 95% CI** | **p_vs0 raw** | **p_vs0 adj** | **Awake PT, Δz mean ± SD (n) 95% CI** | **p_vs0 raw** | **p_vs0 adj** | **Iso PT, Δz mean ± SD (n) 95% CI** | **p_vs0 raw** | **p_vs0 adj** | **Welch F** | **df1** | **df2** | **p_group raw** | **p_group adj** |
| --- | --- | --- | --- | --- | --- | --- | --- | --- | --- | --- | --- | --- | --- | --- | --- |
| Pixelwise healthy hemisphere | 1.5 h − Baseline | 0.138 ± 0.184 (5) 95% CI −0.091…0.366 | 0.170 | 0.509 | −0.019 ± 0.164 (5) 95% CI −0.222…0.185 | 0.813 | 1.000 | 0.028 ± 0.110 (7) 95% CI −0.073…0.130 | 0.519 | 1.000 | 0.97 | 2 | 7.47 | 0.422 | — |
|  | Day 1 − Baseline | 0.070 ± 0.127 (5) 95% CI −0.088…0.228 | 0.286 | 0.572 | 0.210 ± 0.143 (5) 95% CI 0.032…0.388 | **0.031** | **0.092** | 0.057 ± 0.177 (5) 95% CI −0.162…0.277 | 0.509 | 0.572 | 1.55 | 2 | 7.87 | 0.271 | — |
|  | Day 3 − Day 1 | −0.006 ± 0.112 (4) 95% CI −0.184…0.172 | 0.918 | 1.000 | −0.138 ± 0.190 (5) 95% CI −0.373…0.098 | 0.180 | 0.540 | −0.029 ± 0.283 (5) 95% CI −0.380…0.322 | 0.831 | 1.000 | 0.77 | 2 | 7.06 | 0.498 | — |
|  | Day 3 − Baseline | 0.106 ± 0.102 (5) 95% CI −0.021…0.233 | **0.081** | 0.243 | 0.072 ± 0.161 (5) 95% CI −0.127…0.272 | 0.371 | 0.742 | 0.029 ± 0.199 (5) 95% CI −0.219…0.276 | 0.764 | 0.764 | 0.30 | 2 | 7.39 | 0.751 | — |
| MOp | 1.5 h − Baseline | 0.133 ± 0.185 (5) 95% CI −0.096…0.363 | 0.181 | 0.362 | −0.014 ± 0.174 (5) 95% CI −0.230…0.203 | 0.869 | 0.869 | 0.109 ± 0.119 (7) 95% CI −0.001…0.219 | **0.051** | 0.154 | 1.02 | 2 | 7.61 | 0.403 | 1.000 |
|  | Day 1 − Baseline | 0.065 ± 0.147 (5) 95% CI −0.117…0.247 | 0.379 | 0.430 | 0.227 ± 0.207 (5) 95% CI −0.030…0.485 | **0.070** | 0.211 | 0.123 ± 0.187 (5) 95% CI −0.109…0.356 | 0.215 | 0.430 | 0.94 | 2 | 7.82 | 0.430 | 1.000 |
|  | Day 3 − Day 1 | 0.045 ± 0.130 (4) 95% CI −0.162…0.253 | 0.537 | 1.000 | −0.168 ± 0.239 (5) 95% CI −0.466…0.129 | 0.191 | 0.573 | −0.054 ± 0.306 (5) 95% CI −0.434…0.325 | 0.711 | 1.000 | 1.36 | 2 | 7.05 | 0.316 | 0.947 |
|  | Day 3 − Baseline | 0.169 ± 0.111 (5) 95% CI 0.031…0.308 | **0.027** | **0.082** | 0.059 ± 0.179 (5) 95% CI −0.163…0.281 | 0.502 | 0.976 | 0.069 ± 0.202 (5) 95% CI −0.182…0.319 | 0.488 | 0.976 | 0.86 | 2 | 7.44 | 0.462 | 1.000 |
| MOs | 1.5 h − Baseline | 0.113 ± 0.158 (5) 95% CI −0.083…0.309 | 0.185 | 0.556 | −0.017 ± 0.180 (5) 95% CI −0.241…0.206 | 0.840 | 0.840 | 0.043 ± 0.117 (7) 95% CI −0.066…0.151 | 0.375 | 0.749 | 0.70 | 2 | 7.75 | 0.527 | 1.000 |
|  | Day 1 − Baseline | 0.095 ± 0.118 (5) 95% CI −0.052…0.241 | 0.147 | 0.178 | 0.213 ± 0.120 (5) 95% CI 0.063…0.362 | **0.017** | **0.051** | 0.102 ± 0.102 (5) 95% CI −0.025…0.228 | **0.089** | 0.178 | 1.47 | 2 | 7.95 | 0.286 | 1.000 |
|  | Day 3 − Day 1 | −0.069 ± 0.106 (4) 95% CI −0.237…0.099 | 0.280 | 0.560 | −0.122 ± 0.159 (5) 95% CI −0.320…0.076 | 0.162 | 0.486 | −0.022 ± 0.250 (5) 95% CI −0.332…0.288 | 0.856 | 0.856 | 0.30 | 2 | 7.14 | 0.748 | 0.947 |
|  | Day 3 − Baseline | 0.059 ± 0.080 (5) 95% CI −0.040…0.158 | 0.174 | 0.523 | 0.091 ± 0.183 (5) 95% CI −0.136…0.317 | 0.330 | 0.659 | 0.080 ± 0.182 (5) 95% CI −0.146…0.306 | 0.382 | 0.659 | 0.07 | 2 | 6.83 | 0.931 | 1.000 |
| SS | 1.5 h − Baseline | 0.185 ± 0.226 (5) 95% CI −0.096…0.465 | 0.142 | 0.425 | −0.025 ± 0.196 (5) 95% CI −0.269…0.218 | 0.788 | 1.000 | 0.033 ± 0.158 (7) 95% CI −0.113…0.179 | 0.601 | 1.000 | 1.19 | 2 | 7.96 | 0.352 | 1.000 |
|  | Day 1 − Baseline | 0.075 ± 0.158 (5) 95% CI −0.122…0.271 | 0.350 | 0.701 | 0.247 ± 0.267 (5) 95% CI −0.084…0.578 | 0.107 | 0.320 | 0.078 ± 0.185 (5) 95% CI −0.151…0.307 | 0.397 | 0.701 | 0.79 | 2 | 7.72 | 0.488 | 1.000 |
|  | Day 3 − Day 1 | 0.085 ± 0.220 (4) 95% CI −0.265…0.435 | 0.494 | 0.988 | −0.202 ± 0.247 (5) 95% CI −0.508…0.105 | 0.142 | 0.425 | 0.026 ± 0.104 (4) 95% CI −0.140…0.192 | 0.652 | 0.988 | 1.88 | 2 | 5.94 | 0.233 | 0.931 |
|  | Day 3 − Baseline | 0.209 ± 0.183 (5) 95% CI −0.018…0.436 | **0.063** | 0.188 | 0.046 ± 0.149 (5) 95% CI −0.140…0.231 | 0.532 | 0.956 | 0.079 ± 0.197 (4) 95% CI −0.233…0.392 | 0.478 | 0.956 | 1.13 | 2 | 6.79 | 0.377 | 1.000 |
| RSP | 1.5 h − Baseline | 0.216 ± 0.259 (4) 95% CI −0.197…0.629 | 0.195 | 0.584 | −0.047 ± 0.121 (5) 95% CI −0.198…0.104 | 0.438 | 0.875 | −0.038 ± 0.193 (7) 95% CI −0.217…0.140 | 0.620 | 0.875 | 1.64 | 2 | 6.83 | 0.261 | 1.000 |
|  | Day 1 − Baseline | −0.042 ± 0.170 (4) 95% CI −0.312…0.228 | 0.653 | 1.000 | 0.096 ± 0.224 (5) 95% CI −0.183…0.374 | 0.393 | 1.000 | −0.102 ± 0.322 (5) 95% CI −0.502…0.298 | 0.517 | 1.000 | 0.76 | 2 | 7.25 | 0.501 | 1.000 |
|  | Day 3 − Day 1 | 0.050 ± 0.047 (3) 95% CI −0.065…0.166 | 0.202 | 0.508 | −0.083 ± 0.254 (5) 95% CI −0.398…0.233 | 0.507 | 0.508 | 0.140 ± 0.155 (4) 95% CI −0.107…0.387 | 0.169 | 0.508 | 1.19 | 2 | 5.26 | 0.374 | 0.947 |
|  | Day 3 − Baseline | 0.058 ± 0.250 (4) 95% CI −0.339…0.455 | 0.674 | 1.000 | 0.013 ± 0.172 (5) 95% CI −0.200…0.227 | 0.872 | 1.000 | −0.018 ± 0.215 (4) 95% CI −0.361…0.324 | 0.875 | 1.000 | 0.10 | 2 | 5.98 | 0.909 | 1.000 |

**Note.** All changes are Fisher Δz. Pixelwise GBC is calculated for the morphologically intact hemisphere. Contrasts: 1.5 h − baseline; day 1 − baseline; day 3 − day 1; day 3 − baseline. p_vs0 adj, Holm correction across the three experimental groups for one-sample comparisons with zero. For regional rows, p_group adj is Holm-corrected across the four ROIs; the pixelwise scalar has no regional multiplicity correction (—). Values with p < 0.10 are bold burgundy.

**Table S4A. Descriptive Statistics for Motor Outcomes and Serum Biomarkers**

| **Measure** | **Timepoint** | **Laser, mean ± SD (n)** | **Awake PT, mean ± SD (n)** | **Iso PT, mean ± SD (n)** |
| --- | --- | --- | --- | --- |
| Grid test — left-limb errors, n per 100 steps | Baseline | 1.8 ± 1.2 (6) | 1.6 ± 0.5 (5) | 1.4 ± 0.9 (5) |
|  | Day 1 | 9.2 ± 2.1 (6) | 8.4 ± 1.9 (5) | 13.1 ± 4.9 (4) |
|  | Day 3 | 7.4 ± 2.5 (5) | 8.2 ± 3.6 (5) | 13.8 ± 7.0 (4) |
| Cylinder test — asymmetry, % | Baseline | 47.8 ± 1.0 (6) | 49.2 ± 0.8 (5) | 48.6 ± 2.3 (5) |
|  | Day 1 | 44.7 ± 5.5 (6) | 36.2 ± 21.4 (5) | 16.2 ± 15.8 (5) |
|  | Day 3 | 46.2 ± 2.3 (5) | 48.2 ± 9.1 (5) | 28.8 ± 27.0 (5) |
| NSE, pg/mL | Baseline | 200.2 ± 148.3 (5) | 341.8 ± 317.1 (4) | 293.2 ± 102.6 (4) |
|  | Day 1 | 577.9 ± 342.0 (5) | 1499.4 ± 553.3 (3) | 1640.8 ± 558.7 (4) |
|  | Day 3 | 417.9 ± 212.6 (5) | 1237.0 ± 256.7 (5) | 2475.3 ± 682.9 (5) |
| S100b, pg/mL | Baseline | 91.3 ± 76.1 (5) | 26.0 ± 23.2 (3) | 83.8 ± 67.4 (5) |
|  | Day 1 | 34.4 ± 31.6 (3) | 24.6 ± 8.4 (2) | 45.9 ± 47.3 (4) |
|  | Day 3 | 44.1 ± 14.3 (3) | 48.2 ± 34.5 (4) | 37.7 ± 19.2 (5) |

**Table S4B. Longitudinal Changes in Motor Outcomes and Serum Biomarkers: Within-Group Changes from Zero and Between-Group Welch's ANOVA**

| **Measure / contrast** | **Laser, Δ mean ± SD (n) 95% CI** | **p_vs0 raw** | **p_vs0 adj** | **Awake PT, Δ mean ± SD (n) 95% CI** | **p_vs0 raw** | **p_vs0 adj** | **Iso PT, Δ mean ± SD (n) 95% CI** | **p_vs0 raw** | **p_vs0 adj** | **Welch F** | **df1** | **df2** | **p_group raw** | **p_group adj** |
| --- | --- | --- | --- | --- | --- | --- | --- | --- | --- | --- | --- | --- | --- | --- |
| Grid test — left-limb errors Day 1 − Baseline | 7.3 ± 1.8 (6) 95% CI 5.5…9.2 | **<0.001** | **<0.001** | 6.8 ± 2.2 (5) 95% CI 4.1…9.5 | **0.002** | **0.004** | 11.9 ± 5.5 (4) 95% CI 3.2…20.6 | **0.023** | **0.023** | 1.37 | 2 | 6.00 | 0.323 | — |
| Grid test — left-limb errors Day 3 − Day 1 | −1.2 ± 1.8 (5) 95% CI −3.4…1.0 | 0.208 | 0.624 | −0.2 ± 2.9 (5) 95% CI −3.9…3.5 | 0.887 | 1.000 | −0.7 ± 2.1 (3) 95% CI −5.8…4.5 | 0.635 | 1.000 | 0.20 | 2 | 5.40 | 0.822 | — |
| Grid test — left-limb errors Day 3 − Baseline | 5.6 ± 1.8 (5) 95% CI 3.3…7.9 | **0.002** | **0.007** | 6.6 ± 3.4 (5) 95% CI 2.4…10.8 | **0.012** | **0.024** | 12.0 ± 6.8 (4) 95% CI 1.2…22.8 | **0.038** | **0.038** | 1.57 | 2 | 5.60 | 0.287 | — |
| Cylinder test — asymmetry Day 1 − Baseline | −3.2 ± 4.7 (6) 95% CI −8.1…1.7 | 0.157 | 0.315 | −13.0 ± 22.0 (5) 95% CI −40.3…14.3 | 0.256 | 0.315 | −32.4 ± 16.4 (5) 95% CI −52.8…−12.0 | **0.012** | **0.035** | 6.97 | 2 | 5.85 | **0.028** | — |
| Cylinder test — asymmetry Day 3 − Day 1 | 0.8 ± 4.3 (5) 95% CI −4.5…6.1 | 0.697 | 0.697 | 12.0 ± 21.2 (5) 95% CI −14.3…38.3 | 0.273 | 0.547 | 12.6 ± 12.2 (5) 95% CI −2.5…27.7 | **0.081** | 0.244 | 2.37 | 2 | 6.09 | 0.174 | — |
| Cylinder test — asymmetry Day 3 − Baseline | −1.6 ± 1.7 (5) 95% CI −3.7…0.5 | **0.099** | 0.298 | −1.0 ± 8.8 (5) 95% CI −12.0…10.0 | 0.813 | 0.813 | −19.8 ± 27.0 (5) 95% CI −53.3…13.7 | 0.177 | 0.353 | 1.02 | 2 | 5.54 | 0.419 | — |
| NSE, pg/mL Day 1 − Baseline | 355.4 ± 248.2 (4) 95% CI −39.6…750.3 | **0.064** | 0.129 | 1107.3 ± 281.7 (3) 95% CI 407.4…1807.1 | **0.021** | **0.063** | 1375.3 ± 649.5 (3) 95% CI −238.1…2988.7 | **0.067** | 0.129 | 7.30 | 2 | 3.77 | **0.051** | — |
| NSE, pg/mL Day 3 − Day 1 | −27.6 ± 359.8 (4) 95% CI −600.1…544.8 | 0.888 | 0.984 | −275.2 ± 571.2 (3) 95% CI −1694.2…1143.8 | 0.492 | 0.984 | 723.4 ± 1118.9 (4) 95% CI −1057.0…2503.8 | 0.287 | 0.860 | 1.02 | 2 | 4.28 | 0.433 | — |
| NSE, pg/mL Day 3 − Baseline | 256.1 ± 317.0 (4) 95% CI −248.3…760.4 | 0.205 | 0.205 | 869.2 ± 247.8 (4) 95% CI 474.9…1263.4 | **0.006** | **0.018** | 2245.2 ± 732.9 (4) 95% CI 1079.0…3411.3 | **0.009** | **0.018** | 12.16 | 2 | 5.46 | **0.010** | — |
| S100b Day 1 − Baseline | −96.8 ± 104.4 (3) 95% CI −356.1…162.4 | 0.249 | 0.430 | 0.0 (n=1; SD NA) 95% CI NA | NA | NA | −48.3 ± 61.6 (4) 95% CI −146.4…49.8 | 0.215 | 0.430 | NA | 2 | NA | NA | — |
| S100b Day 3 − Day 1 | 35.9 (n=1; SD NA) 95% CI NA | NA | NA | −5.3 (n=1; SD NA) 95% CI NA | NA | NA | −10.9 ± 66.8 (4) 95% CI −117.3…95.4 | 0.765 | 0.765 | NA | 2 | NA | NA | — |
| S100b Day 3 − Baseline | −76.3 ± 123.8 (2) 95% CI −1188.4…1035.9 | 0.544 | 0.562 | 40.4 ± 10.5 (2) 95% CI −53.9…134.7 | 0.116 | 0.347 | −46.1 ± 82.8 (5) 95% CI −148.9…56.7 | 0.281 | 0.562 | 2.65 | 2 | 2.19 | 0.260 | — |

**Note.** Contrasts: day 1 − baseline; day 3 − day 1; day 3 − baseline. p_vs0 adj, Holm correction across the three groups for one-sample comparisons with zero. Between-group Welch ANOVA is a scalar test and has no additional regional correction (p_group adj = —). NA denotes statistics not estimable from the available paired sample. Values with p < 0.10 are bold burgundy.

**Table S5A. OISI-Derived Lesion Area: Descriptive Statistics and Between-Group Welch's ANOVA**

| **Timepoint** | **Laser, mean ± SD (n), mm²** | **Awake PT, mean ± SD (n), mm²** | **Iso PT, mean ± SD (n), mm²** | **Welch F** | **df1** | **df2** | **p_group** |
| --- | --- | --- | --- | --- | --- | --- | --- |
| 10 min | 2,03 ± 0,99 (6) | 3,54 ± 1,58 (5) | 6,24 ± 1,36 (7) | 19,40 | 2 | 8,94 | **<0,001** |
| 1.5 h | 4,41 ± 1,16 (6) | 6,84 ± 1,46 (5) | 8,00 ± 3,03 (7) | 6,65 | 2 | 9,28 | **0,016** |
| Day 1 | 5,60 ± 1,38 (6) | 9,31 ± 1,49 (5) | 10,04 ± 2,57 (5) | 10,89 | 2 | 7,83 | **0,005** |
| Day 3 | 4,07 ± 1,16 (5) | 9,29 ± 3,00 (5) | 12,80 ± 1,87 (5) | 37,92 | 2 | 7,12 | **<0,001** |

**Note.** Lesion area was manually delineated from OISI data. Absolute lesion areas are shown for 10 min, 1.5 h, day 1, and day 3 after injury. Values with p < 0.10 are bold burgundy.

**Table S5B. Pairwise Between-Group Comparisons of OISI-Derived Lesion Area**

| **Timepoint** | **Comparison** | **Difference, mm²** | **95% CI** | **t** | **df** | **p_raw** | **p_adj** | **Hedges g** |
| --- | --- | --- | --- | --- | --- | --- | --- | --- |
| 10 min | Awake PT − Laser | +1.52 | −0.44…3.48 | 1.86 | 6.46 | 0.109 | 0.109 | +1.08 |
| 10 min | Iso PT − Laser | +4.22 | 2.77…5.66 | 6.46 | 10.76 | **<0.001** | **<0.001** | +3.26 |
| 10 min | Iso PT − Awake PT | +2.70 | 0.68…4.72 | 3.08 | 7.87 | **0.015** | **0.031** | +1.71 |
| 1.5 h | Awake PT − Laser | +2.43 | 0.56…4.30 | 3.01 | 7.65 | **0.018** | **0.053** | +1.71 |
| 1.5 h | Iso PT − Laser | +3.59 | 0.73…6.46 | 2.89 | 7.95 | **0.020** | **0.053** | +1.41 |
| 1.5 h | Iso PT − Awake PT | +1.16 | −1.82…4.14 | 0.88 | 9.08 | 0.401 | 0.401 | +0.43 |
| Day 1 | Awake PT − Laser | +3.71 | 1.71…5.71 | 4.25 | 8.36 | **0.003** | **0.008** | +2.37 |
| Day 1 | Iso PT − Laser | +4.44 | 1.30…7.59 | 3.47 | 5.89 | **0.014** | **0.027** | +2.03 |
| Day 1 | Iso PT − Awake PT | +0.73 | −2.47…3.93 | 0.55 | 6.42 | 0.602 | 0.602 | +0.31 |
| Day 3 | Awake PT − Laser | +5.22 | 1.55…8.88 | 3.63 | 5.16 | **0.014** | **0.029** | +2.07 |
| Day 3 | Iso PT − Laser | +8.73 | 6.39…11.08 | 8.89 | 6.68 | **<0.001** | **<0.001** | +5.08 |
| Day 3 | Iso PT − Awake PT | +3.51 | −0.26…7.29 | 2.22 | 6.69 | **0.063** | **0.063** | +1.27 |

**Note.** Differences are expressed in the direction shown in the Comparison column. p_adj, Holm correction across the three planned biological contrasts within each timepoint. Values with p < 0.10 are bold burgundy; |g| ≥ 0.80 is burgundy.

**Table S5C. Longitudinal Changes in OISI-Derived Lesion Area**

| **Contrast** | **Laser, Δ mean ± SD (n) 95% CI** | **p_vs0 raw** | **p_vs0 adj** | **Awake PT, Δ mean ± SD (n) 95% CI** | **p_vs0 raw** | **p_vs0 adj** | **Iso PT, Δ mean ± SD (n) 95% CI** | **p_vs0 raw** | **p_vs0 adj** | **Welch F** | **df1** | **df2** | **p_group raw** | **p_group adj** |
| --- | --- | --- | --- | --- | --- | --- | --- | --- | --- | --- | --- | --- | --- | --- |
| Day 1 − 1.5 h | 1.19 ± 0.99 (6) 95% CI 0.15…2.23 | **0.032** | **0.097** | 2.47 ± 2.02 (5) 95% CI −0.03…4.97 | **0.052** | 0.104 | 0.66 ± 0.92 (5) 95% CI −0.48…1.80 | 0.185 | 0.185 | 1.62 | 2 | 7.83 | 0.257 | — |
| Day 3 − Day 1 | −1.69 ± 1.53 (5) 95% CI −3.58…0.21 | **0.069** | 0.207 | −0.02 ± 2.38 (5) 95% CI −2.97…2.93 | 0.985 | 0.985 | 2.76 ± 2.69 (5) 95% CI −0.58…6.11 | **0.083** | 0.207 | 4.85 | 2 | 7.49 | **0.045** | — |

**Note.** Changes are in mm². Contrasts: day 1 − 1.5 h and day 3 − day 1. p_vs0 adj, Holm correction across the three groups for one-sample comparisons with zero. Between-group Welch ANOVA has no additional regional correction (p_group adj = —). Values with p < 0.10 are bold burgundy.

**Table S5D. Pairwise Between-Group Comparisons of Longitudinal Changes in OISI-Derived Lesion Area**

| **Contrast** | **Comparison** | **Difference in Δ, mm²** | **95% CI** | **t** | **df** | **p_raw** | **p_adj** | **Hedges g** |
| --- | --- | --- | --- | --- | --- | --- | --- | --- |
| Day 1 − 1.5 h | Awake PT − Laser | +1.28 | −1.18…3.74 | 1.30 | 5.59 | 0.245 | 0.490 | +0.76 |
| Day 1 − 1.5 h | Iso PT − Laser | −0.53 | −1.84…0.77 | −0.92 | 8.86 | 0.380 | 0.490 | −0.51 |
| Day 1 − 1.5 h | Iso PT − Awake PT | −1.82 | −4.28…0.65 | −1.83 | 5.59 | 0.120 | 0.360 | −1.05 |
| Day 3 − Day 1 | Awake PT − Laser | +1.67 | −1.34…4.67 | 1.32 | 6.83 | 0.230 | 0.243 | +0.75 |
| Day 3 − Day 1 | Iso PT − Laser | +4.45 | 1.11…7.80 | 3.21 | 6.33 | **0.017** | **0.051** | +1.84 |
| Day 3 − Day 1 | Iso PT − Awake PT | +2.79 | −0.93…6.50 | 1.73 | 7.88 | 0.122 | 0.243 | +0.99 |

**Note.** Differences are expressed in the direction shown in the Comparison column. p_adj. Holm correction across the three planned biological contrasts within each longitudinal endpoint. Values with p < 0.10 are bold burgundy; |g| ≥ 0.80 is burgundy.
